# Generalizing biological surround suppression based on center surround similarity via deep neural network models

**DOI:** 10.1101/2023.03.18.533295

**Authors:** Xu Pan, Annie DeForge, Odelia Schwartz

## Abstract

Sensory perception is dramatically influenced by the context. Models of contextual neural surround effects in vision have mostly accounted for Primary Visual Cortex (V1) data, via nonlinear computations such as divisive normalization. However, surround effects are not well understood within a hierarchy, for neurons with more complex stimulus selectivity beyond V1. We utilized feedforward deep convolutional neural networks and developed a gradient-based technique to visualize the most suppressive and excitatory surround. We found that deep neural networks exhibited a key signature of surround effects in V1, highlighting center stimuli that visually stand out from the surround and suppressing responses when the surround stimulus is similar to the center. We found that in some neurons, especially in late layers, when the center stimulus was altered, the most suppressive surround surprisingly can follow the change. Through the visualization approach, we generalized previous understanding of surround effects to more complex stimuli, in ways that have not been revealed in visual cortices. In contrast, the suppression based on center surround similarity was not observed in an untrained network. We identified further successes and mismatches of the feedforward CNNs to the biology. Our results provide a testable hypothesis of surround effects in higher visual cortices, and the visualization approach could be adopted in future biological experimental designs.

**Author summary:** Neural responses and perception of a visual stimulus are influenced by the context, such as what spatially surrounds a given feature. Contextual surround effects have been extensively studied in the early visual cortex. But the brain processes visual inputs hierarchically, from simple features up to complex objects in higher visual areas. Contextual effects are not well understood for higher areas of cortex and for more complex stimuli. Utilizing artificial deep neural networks and a visualization technique we developed, we found that deep networks exhibited a key signature of surround effects in the early visual cortex, highlighting center stimuli that visually stand out from the surround and suppressing responses when the surround stimulus is similar to the center. We found in some neurons, especially in late layers, when the center stimulus was altered, the most suppressive surround could surprisingly follow. This is a generalization of known surround effects for more complex stimuli that has not been revealed in the visual cortex. Our findings relate to notions of efficient coding and salience perception, and emerged without incorporating specialized nonlinear computations typically used to explain contextual effects in the early cortex. Our visualization approach provides a new experimental paradigm and a testable hypothesis of surround effects for more complex stimuli in higher cortical areas; the visualization approach could be adopted in biological experimental designs.

## Introduction

Both biological and artificial systems seek to make sense of complex structured information in the world. A key aspect of sensory input is that its interpretation at a given point depends on the context, for example, what surrounds a given feature or object. Spatial context in vision plays a role in perceptual grouping [1] and segmentation [2], highlighting salient objects in which a stimulus stands out from its background [3], and resulting in visual illusions [4, 5]. Deficits have been associated with disorders [6–8]. Though contextual surround effects are ubiquitous in visual cortex, they are not well understood within hierarchical systems such as deep neural networks and for neurons with more complex stimulus selectivity beyond V1.

A rich set of surround effects have been documented in the Primary Visual Cortex (V1) in neurophysiology experiments and respective modeling studies [9–29]. In the experiments, researchers typically place a stimulus in the center (i.e. the classical receptive field) and in the surround (i.e. beyond the classical receptive field). Studies have found that a surround stimulus that does not elicit a response by itself, may nevertheless nonlinearly modulate the response to a center stimulus. Modeling studies have addressed V1 data by incorporating nonlinear computations such as divisive normalization or dynamical circuitry [14, 19, 23, 29, 30].

Surround effects are less well understood in cortical areas beyond V1 (though see [31, 32]). Moreover, surround suppression in V2 for textures versus noise [31] cannot be simply explained by divisive normalization models that have been successful for V1 data. Therefore, novel experimental paradigms and hierarchical models that make predictions on complex features are in demand to study surround effects in higher visual areas. In recent years, Deep Convolutional Neural Networks (CNNs) that stack up multiple layers of computation have achieved astonishing visual task performance and have been used as descriptive models to capture visual neuron properties across the cortical hierarchy [33–41]. But beyond the observation that deep neural networks can exhibit surround suppression [41], it is not clear what properties of the center and surround stimuli lead to surround suppression; to what extent feedforward CNNs that lack specialized nonlinear computations such as divisive normalization and lateral or feedback connections can capture the rich surround effects that have been studied biologically; and excitingly, what predictions CNNs can make about surround effects in higher visual cortex with complex stimuli.

Moreover, feature visualization techniques have become popular in neurophysiology experiments [42–44] and in analyzing what stimuli most excite CNN artificial neurons [45–47]. However, neither in CNN studies nor in neurophysiology, have such techniques been extended to visualizing surround effects. Developing surround visualization techniques could address the limitation in current neurophysiology studies that the surround stimuli are usually simple parametric stimuli or are selected from a fixed set of textures or natural images.

Utilizing feedforward deep neural networks and developing a novel gradient-based visualization technique, we found that CNN neurons exhibit a key signature of surround suppression, namely that on average they are most suppressed when the surround matches the center and less suppressed when the surround differs from the center; and that this can even follow when the center orientation is altered. Suppression based on center surround similarity is known for V1 data [48], but has not been observed in higher cortical areas. These findings generalize the idea of homogeneity-dependent surround suppression to more complex stimuli [26], thus providing a testable hypothesis of surround effects in higher visual cortices. Surround suppression for homogeneous center and surround can highlight center stimuli that stand out from the surround, relating to visual salience [3]. Suppression based on center-surround similarity also relates to notions of efficient coding. Note that we use the term homogeneity to indicate similarity of the center and surround in terms of stimulus features such as orientation, color, spatial frequency and textures, rather than examining the conditions of statistical similarity as in some modeling studies of natural stimuli [26]. The visualization method reveals a generalization of the idea of homogeneity to complex stimuli and provides a new experimental scheme that can be used in biological experiments.

Although neurons in feedforward neural networks do not have a clear separation between the center and surround region, we suggest that at a computational level they nevertheless exhibit some surround effects that are prototypical in primary visual cortex neurons. Our results can partly be attributed to the weight kernel of the CNN neurons that includes positive and negative weights, and from the spatial integration of rectified neurons in the previous layer. While this is not recurrent connections as a mechanism, this serves as including lateral inputs that could be positive or negatively weighted from neurons in the previous layer. This might be a way for feedforward CNNs to achieve some of the noted surround suppression effects observed biologically. In contrast, we did not find such effects in an untrained CNN network, suggesting that the architecture alone was not sufficient and that learning on images was important.

We also found mismatches to the biology, highlighting the limitations of the feedforward architectures, and identifying the need to further incorporate nonlinear computations and circuitry into deep neural networks [49–54].

## Results

### Defining center and surround in CNN neurons

Before studying the surround effects in CNN neurons, we first defined the center and surround region through a method inspired by neurophysiology studies [16] (Fig 1). We used two standard feedforward network architectures, Alexnet [55] and VGG16 [56], which have been applied extensively in neural modeling (see Methods). Without losing generality, we focused on the center neurons in each feature map. However, unlike cortical neurons, each CNN neuron has a well-defined theoretical receptive field (see Methods and Supplementary Figure 1). We used this theoretical receptive field as the outer size of the stimuli in the following experiments. To define a center region for each of the CNN neurons, we adopted a physiology approach [16]: first, we used a grid search to find the optimal spatial frequency and orientation for each neuron; then we grew the size of the optimal stimulus and computed the neural response as a function of the stimulus diameter (i.e., the diameter tuning curve) to find the extent of the center receptive field. For the boundary between the center and surround, we used the grating summation field, defined as the diameter that elicits at least 95% of the peak responses of the diameter tuning curve (see Methods and Figure 1 top). For the grating summation field population distribution, see Supplementary Figure 3. Following the convention in neurophysiology, we defined the suppression index as the fractional reduction in responses to optimal diameter. We found that the suppression index is higher in later layers than in earlier layers (Supplementary Figure 3), which is consistent with a previous study [41].

**Fig 1.**
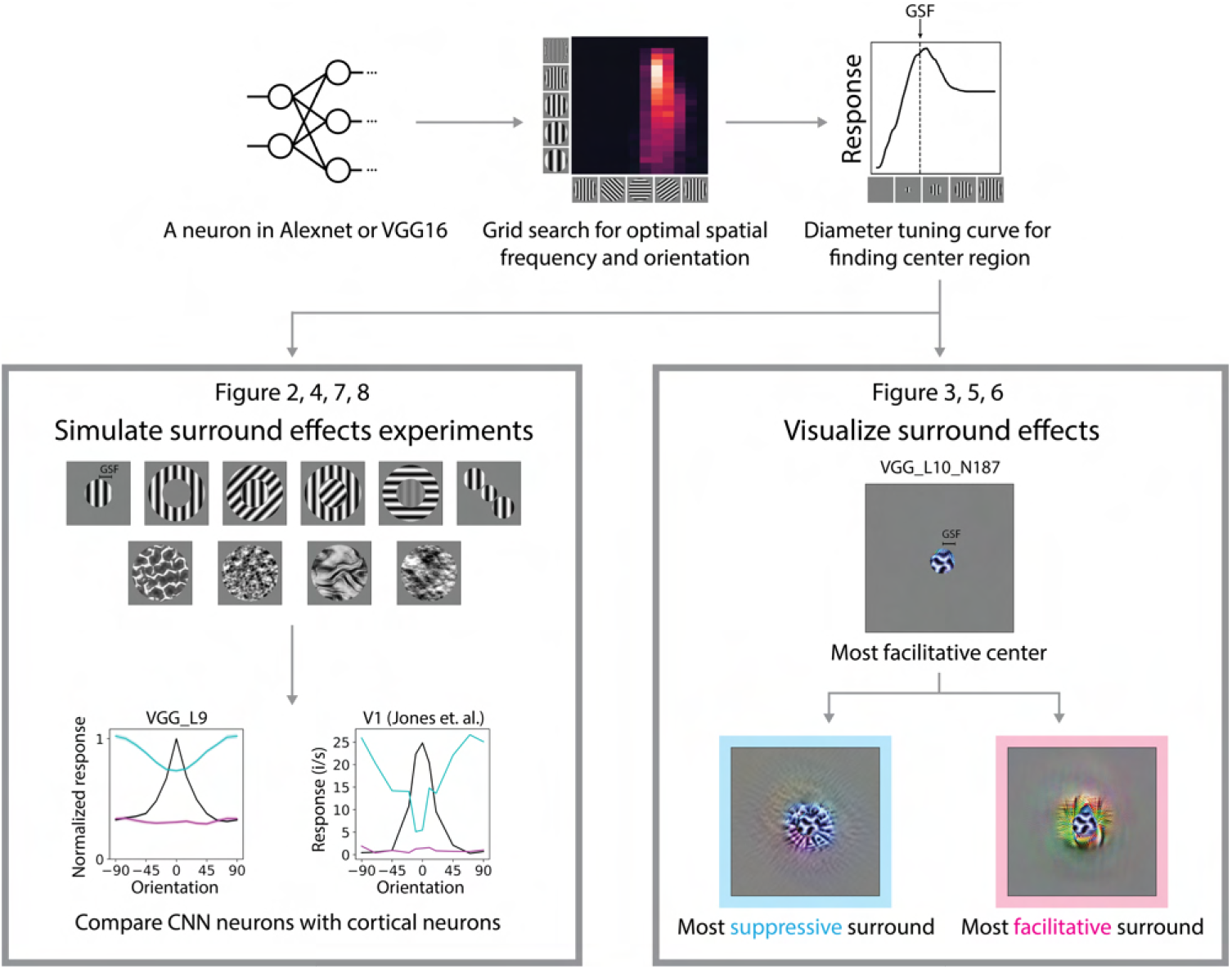
Probing surround effects in CNNs. Top left: A neuron was taken from either Alexnet or VGG16. Top middle: The optimal spatial frequency and grating orientation were found by grid search. Top right: Then the grating summation field (GSF) was read from the grating diameter tuning curve. Bottom left: We simulated a set of in-silico physiology experiments with the stimuli that were used in neurophysiology studies. Representative stimuli are shown. The responses of CNN neurons are compared with cortical neurons. Bottom right: We visualized surround effects in CNN neurons by a two-step optimization approach. First, the most facilitative center was optimized within the grating summation field. Then, the most suppressive and facilitative surround were optimized with the fixed most facilitative center.

By the conventional definition used in neuroscience, a stimulus placed outside the classical receptive field by itself does not elicit any neural responses (Fig 1, Fig 2B) and approaches have been developed to minimize the impingement of the surround onto the center cavanaugh2002. In practice, it is difficult to achieve a complete separation between the center and surround, and center and surround in biological neurons may be considered a continuum [57]. CNN neurons do not have a clear separation of center and surround. The surround orientation tuning curves were on average flat and low (Fig 2D), which makes it a reasonable analogy to the surround region in neurophysiology studies. However, we did find noisy neurons in which the surround alone elicits a response and did not have a clear center surround separation, for example Alexnet L3 N97 in Fig 2C.

**Fig 2.**
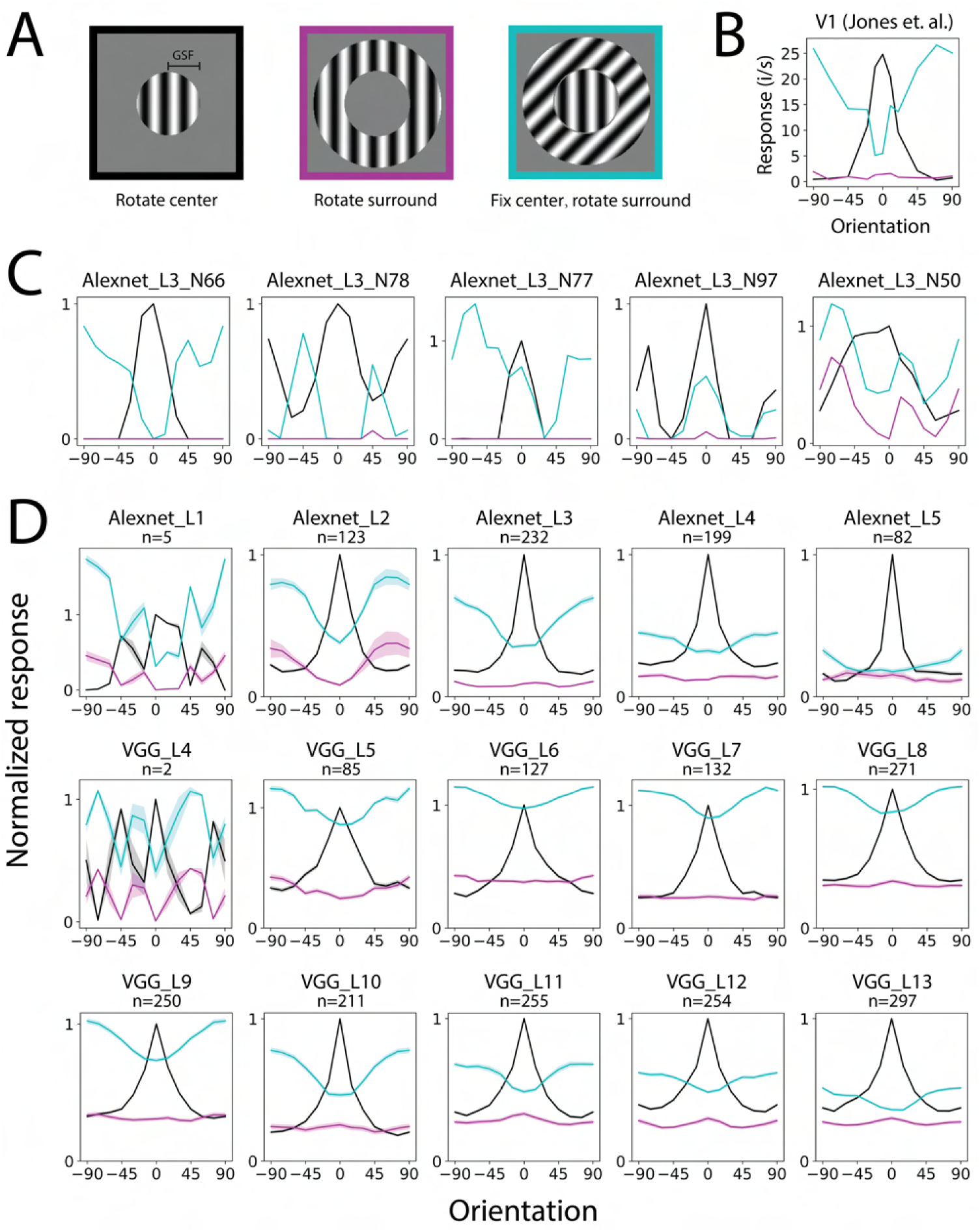
Grating orientation tuning of the CNN neurons. A. Stimuli used in the experiments: rotating the center (left, black); rotating the surround (middle, purple); fixing the center at the optimal orientation and rotating the surround (right, cyan). B. Neurophysiology V1 data of the three types of orientation tuning curves (reproduced from [17]). The most suppressive surround orientation matches the optimal center orientation. The surround stimuli alone hardly elicit responses. 0° represents the optimal orientation (same for the following plots). C. Example orientation tuning curves of CNN neurons. D. Averaged orientation tuning curves in CNN layers. Shaded area indicates s.e.m.

There is a widespread belief that surround effects are due to recurrent and feedback connections, though feedforward mechanisms have also been noted in previous studies [57–63]. Even though the CNN models we studied do not contain recurrent connections explicitly, we think the observed surround effects have connections to their biological counterparts. A computation can be implemented in different ways in different systems. For example, a recurrent neural network can be unrolled in time to a feed-forward neural network [64]. This study is mainly focused on the surround effects at a computational level instead of an implementation level, though we further strive to explain how the effects may arise in the CNN. Though the brain and CNNs are wired in drastically different ways, it is meaningful to compare the computations between the two.

### The most suppressive surround grating matches the optimal orientation

First, we tested one of the most well-known surround effects found in V1 that the surround induces the largest response suppression when the grating orientations of the center and surround are the same [15, 17, 25]. We computed three types of orientation tuning curves: the center orientation tuning curve, the surround orientation tuning curve, and the surround suppression orientation tuning curve for stimuli with a fixed optimal center and a varying surround orientation (surround suppression tuning curve for abbreviation) (Fig 2A). We only included neurons with sufficiently large center and surround (grating summation field in between 30% and 70% of the theoretical receptive field) and the center orientation turning or surround suppression curve has more than 0.001 variation (not silent) in the analysis. The exact number of neurons excluded by each of these criteria is shown in supplementary Table 1. Due to the tiny receptive fields in the early layers in VGG16, only layer 4 and successors had neurons that satisfied these criteria. We sanity-checked the selectivity of orientation in CNN neurons (Supplementary Figure 3) and found it qualitatively consistent with a previous study [41, 65].

In neurophysiology studies, the most suppressive surround has the same orientation as the optimal center orientation, and the surround can be facilitative when it differs from the center [15, 17, 25] (Fig 2B). We found that on average, most layers in both CNNs showed the most suppression when the surround orientation matched the center and the least suppression (and even facilitation in some layers of VGG16) when the orientations differed. This similarity between the CNNs and the neurophysiology held for most layers, except for Alexnet layer 1 and VGG16 layer 4 which lacked sufficient neurons due to the small receptive fields and not meeting our selection criteria (Fig 2D). We further found that when the center contrast is low, there is less influence of whether the surround orientation matches the center or is orthogonal to the center (Supplementary Fig 2). Such finding aligns with neurophysiology studies [9, 10, 15, 21, 66].

We found that the surround suppression in the early layers was weaker than in the late layers. Regarding the amount of surround suppression, Alexnet layer 3 and VGG16 layer 10 were closest to the V1 data. In the neurophysiology data, the strongest responses in the center tuning curve are aligned with the strongest suppression in the surround suppression tuning curve. To examine this quantitatively in the CNN, we measured the negative correlation between the two curves. Consistent with the neurophysiology observations (-0.858 in [17]), all layers of the CNN showed significant negative mean correlations between the two curves (Supplementary Fig3-5).

Note that the averaged tuning curves in Fig 2D did not directly imply the orientation selectivity. The relationship between the surround suppression tuning curve and center orientation tuning curve is what is actually informative. The center orientation tuning curves appeared to have a peak even in late layers. This did not imply those neurons were highly tuned to the oriented gratings. This was just an artifact of aligning the optimal orientation at 0 degrees. Orientation selectivity was actually low in most layers (Supplementary Figure 3) as in previous reports [41, 65]. As an example, averaging multiple samples of random noise can result in a peak when the maximum is aligned. Untrained networks also exhibited a peak in the center orientation tuning curves, but their surround suppression tuning curves did not show any relation to the center orientation tuning curves and thus appeared flat (Supplementary Fig16).

By screening individual neurons, we found that there were a variety of interesting surround suppression behaviors that had not been documented in neurophysiology studies (Fig 2C). This included neurons with a double-peak center orientation tuning curve, for which their surround suppression curve matched both peaks (Alexnet L3 N78); neurons for which their most suppressive surround orientation did not match the center orientation (Alexnet L3 N77); and neurons for which their surround suppression curve matched the center orientation tuning curve (Alexnet L3 N97). We also found neurons that had overall noise curves in which the surround alone elicited responses and the surround suppression curve appeared not related to the center orientation tuning curve(Alexnet L3 N50).

In this and the following analysis, we wanted to use only the few necessary neuron selection criteria to show the population results of as many neurons as possible. On average, the population showed the effects, and there were individual neurons that showed clean effects. However, there were also many neurons with noisy tuning curves that showed less/no effect, like the neuron Alexnet L3 N97 in Fig 2C.

### Visualization of the most suppressive surround appears homogeneous to the center

In both neuroscience and machine learning, there is interest in understanding what visual features neurons are sensitive to. Indeed, with recent advances in deep neural networks, there has been some focus on visualizing what input features induce the most or the least responses in CNN neurons, for instance using gradient based optimization methods [45]. Inspired by neurophysiology studies, we were interested in going beyond such methods and visualizing the most suppressive and facilitative surround and testing if the homogeneous surround induces the most suppression is still applicable to complex stimuli that are beyond gratings. We therefore modified the gradient-based optimization approach to a two-step optimization: first, we optimized the stimuli inside the center region to elicit the strongest response; then, we optimized the stimuli in the surround region that suppressed or facilitated the strongest response when combined with the optimal center stimuli (Fig 1) (see Methods). An advantage of optimizing via two steps over one step is that we can separate the center and surround components more clearly; thus we can study the questions such as what are the most suppressive surround when the center is not optimal.

Figure 3 shows a curation of the visualizations (see the full set in the online repository https://gin.g-node.org/xupan/CNN_surround_effects_visualization). We selected them to show the variety. In general, the most suppressive surround looked similar to the center, whereas the most facilitative surround looked dissimilar to the center. Based on the visual appearance, we found several typical patterns. We observed visual similarity along various features, such as color and spatial frequency; the most suppressive surround could have similar color or spatial frequency to the center (Fig 3C). Visualizations in untrained CNNs did not show textural patterns, but appeared as spectrally matched noise (since the optimization was parameterized to capture the natural image spectrum), and thus did not show these effects (Supplementary Figure 16).

**Fig 3.**
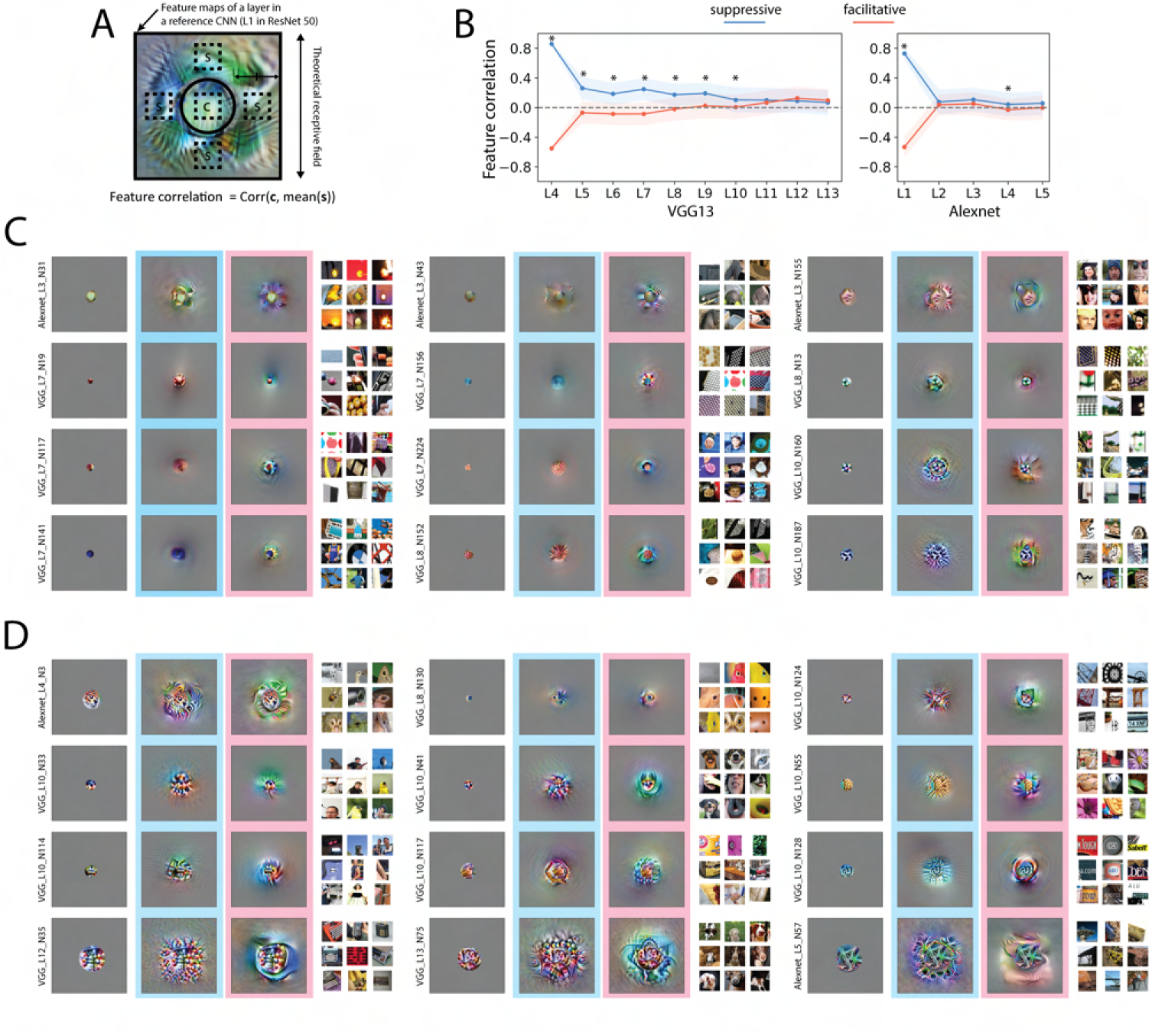
Visualizing the most suppressive and facilitative surround. A. Center surround similarity can be quantified by the feature correlation, in which we took a feature map in another reference CNN, in this case, layer 1 in ResNet50, and computed the correlation between the center features and the average of the surround features. More specifically, four locations (top, bottom, left, right) in the reference feature map that were closest to the middle between the gsf and the theoretical receptive field were used to get surround feature vectors. Feature correlation was calculated as the correlation coefficient between the mean of the four surround feature vectors and the center feature vector. B. Feature correlations in two CNNs. The shaded area indicates standard deviation. Asterisks indicate p value smaller than 0.05 in paired t-test. Note that the feature correlation depends on the selection of reference CNN. Feature correlations calculated with other reference CNNs can be found in Supplementary Figure 6. C, D. The most facilitative center (left image with no frame), most suppressive surround (middle image with cyan frames), and most facilitative surround (right image with pink frames) are shown for each selected neuron. The nine most excitatory natural image patches from the ImageNet validation set are shown on the right for each neuron. C. Example neurons in early layers that have recognizable features: color (left column) and frequency (middle and right column). The most suppressive surrounds appeared similar to the center, whereas the most facilitative surrounds appeared different from the center. D. Example neurons in late layers that have more complex patterns.

We used two metrics to quantify the visual similarity between the center and surround. First, we quantified the color similarity between the center and surround, by calculating the Pearson correlation coefficient between the 3 averaged color channels in the center and surround (Fig 5A and Supplementary Figure 7) as a metric. The correlation values, therefore, ranged between -1 and 1, with a correlation value of 1 indicating that the center and surround have the same color. The most suppressive surround showed a high (positive) color correlation with the center, whereas the most facilitative surround showed a low (negative) color correlation in all layers (Fig 5A and Supplementary Figure 7). We further quantified the similarity between the center and surround based on more general features from a feature map in a reference CNN and used again the Pearson correlation coefficient between the feature vectors of the center and surround as a measure of homogeneous, i.e. feature correlation (Figure 3A, B and Supplementary Figure 6, 7). Note that the feature correlation depends on the selection of the reference feature map, for which several are shown in Supplementary Figure 6 in addition to the one shown in Figure 3B. Significant different feature correlations between the case of the suppressive and facilitative visualization are consistently found in early layers in VGG16 with different reference feature maps (Figure 3B and Supplementary Figure 6). The population distribution of color correlation and feature correlation are shown in Supplementary Figure 7. Our two metrics indeed showed that in most layers, the most suppressive surround was more similar to the center than the most facilitative surround.

Many neurons showed combined features of color and spatial frequency. And in deeper layers, the visual similarity between the center and the most suppressive surround could be more complex (Fig 3D). For example, the color similarity was not limited to a single color, but to a color scheme (VGG L10 N124, VGG L10 N33, VGG L10 N114, etc.); if a swirl was in the center, the most suppressive surround could include several swirls (Alexnet L4 N3, VGG L8 N130, VGG 133 N75, etc.); the line shapes of the center and most suppressive surround matched (VGG L10 N124, VGG L10 N114, VGG L10 128, Alexnet L5 N57, etc.).

These effects were not rare in the CNN neurons; we found that most neurons showed visual similarity/dissimilarity between the most suppressive/facilitative surround and the center to some extent. For a full visualization of all neurons in the two CNNs, see the online repository. However, we found neurons that did not show this effect, especially when the surround features were geometrically arranged rather than uniform across the surround, and when the features were arranged as object-like shapes (Supplementary Fig 8).

Our visualizations align with findings from neurophysiology studies that the most suppressive surround occurs when the center and surround are homogeneous [15, 17, 26, 48], but go beyond simple stimuli and early processing stages.

### The most suppressive grating surround follows the change of the center orientation

An interesting interaction between the center and surround observed in neurophysiology studies is that even when the center grating orientation is not optimal, the most suppressive surround orientation still matches the non-optimal center orientation. This has been documented in V1 neurons [15, 48] (Fig 4B). We tested this effect in the CNN neurons. We used a similar design to the neurophysiology studies [15], setting the center orientation at 0°, 15°, 30°, and 45° degrees off from the optimal orientation, and rotating the surround. We obtained four surround suppression curves for four center orientations and each neuron (Fig 4A). On average, later layers (Layer 6 and successors) in VGG16 captured this effect, with shifted dips matching the center orientation (Fig 4D). This trend was less pronounced in Alexnet. This effect also can be seen in the population histograms of the deviation of the most suppressive surround orientation from the center orientation (Supplementary Figure 9). It is surprising that CNNs could capture this effect to some extent, since all previous successful models of surround effects included non-linear interactions between the center and surround (e.g., in a divisive manner). It appears that even without an explicitly divisive surround, CNNs could still achieve similar center-surround interactions by stacking layers. However, we did not see this effect in more shallow networks and early layers of deep networks (e.g. a 5-layer Alexnet, and earlier layers of VGG16), indicating the computations may not be complex enough to support this interaction (Fig 4D).

**Fig 4.**
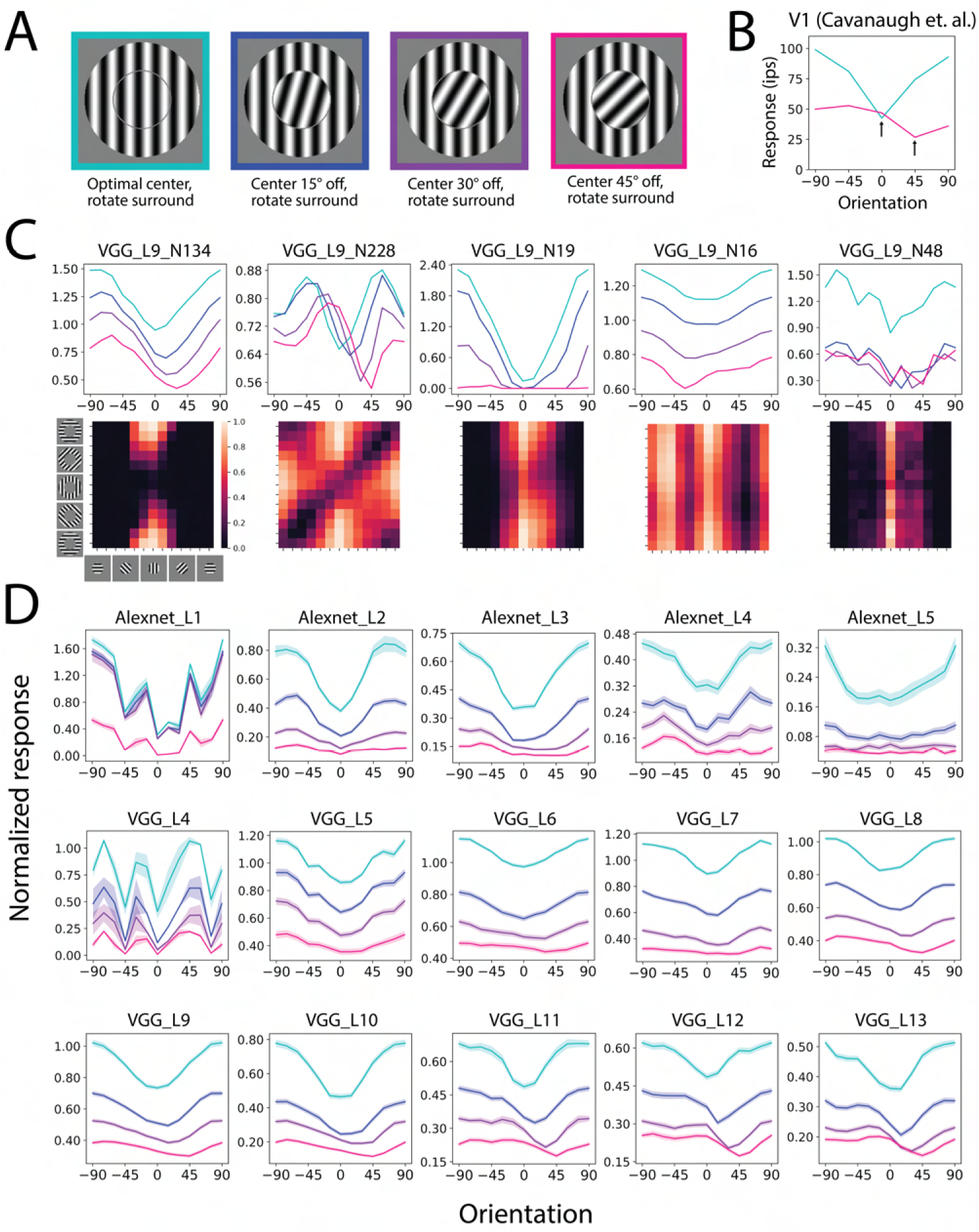
Surround suppression tuning when the center is not at the optimal orientation. A. Stimuli used in the experiments: the center was either fixed at the optimal orientation or rotated 15°, 30°, 45° off from the optimal orientation. The surround suppression tuning curve was acquired by changing the surround orientation. B. Neurophysiology V1 data of Surround suppression tuning curves, when the center was either optimal or rotated 45° away from the optimal (reproduced from [15]). Arrow indicates the center orientation. The most suppressive surround matched the center orientation. C. First row: example surround suppression tuning curves of CNN neurons. We chose these neurons to show the variety of behavior. Second row: Activation heat map of an extended experiment that used more surround and center orientation combinations. Low activation on the diagonal line indicates that the most suppressive surround orientations can follow the center. D. Averaged surround suppression tuning curves in CNN layers. Shaded area indicates s.e.m.

Although the average effects were consistent with the biology, we also found a variety of untypical behaviors (Fig 4C). When the curve had two peaks/dips, some neurons showed a shift of both dips (VGG L9 N228). Some neurons also showed a uniform drop of the curves without shifting the dips (VGG L9 N19). Interestingly, some neurons showed dip shifts in the opposite direction (VGG L9 N16). They may play a role in completing the representation space. We are not aware if such untypical behaviors have been found in the brain.

### Visualization of the most suppressive surround follows changes in the center

Since in the above simulation the homogeneity idea was observed for a non-optimal center grating orientation, we asked if such effects can be revealed in visualizations and generalized for complex stimuli. We altered the optimal center stimuli in two ways and tested if the most suppressive surround can follow the change in the center. First, for each neuron we permuted the three color channels, i.e. red, green, and blue, of the center stimuli. Then we computed the most suppressive and facilitative surround as before (Fig 5). We found that for many neurons the most suppressive surround matched the altered center color. The averaged color correlations are shown in Figure 5A. The altered center of most later layers (after layer 5 in VGG16) had positive color correlations with the most suppressive surround and negative color correlations with the most facilitative surround, though the magnitudes of correlation/anticorrelation were smaller than the optimal center. The color correlation distribution is shown in Supplementary Figure 7. This effect was less pronounced in Alexnet, which indicates the CNNs may need a sufficient number of layers to achieve this type of nonlinear effect.

**Fig 5.**
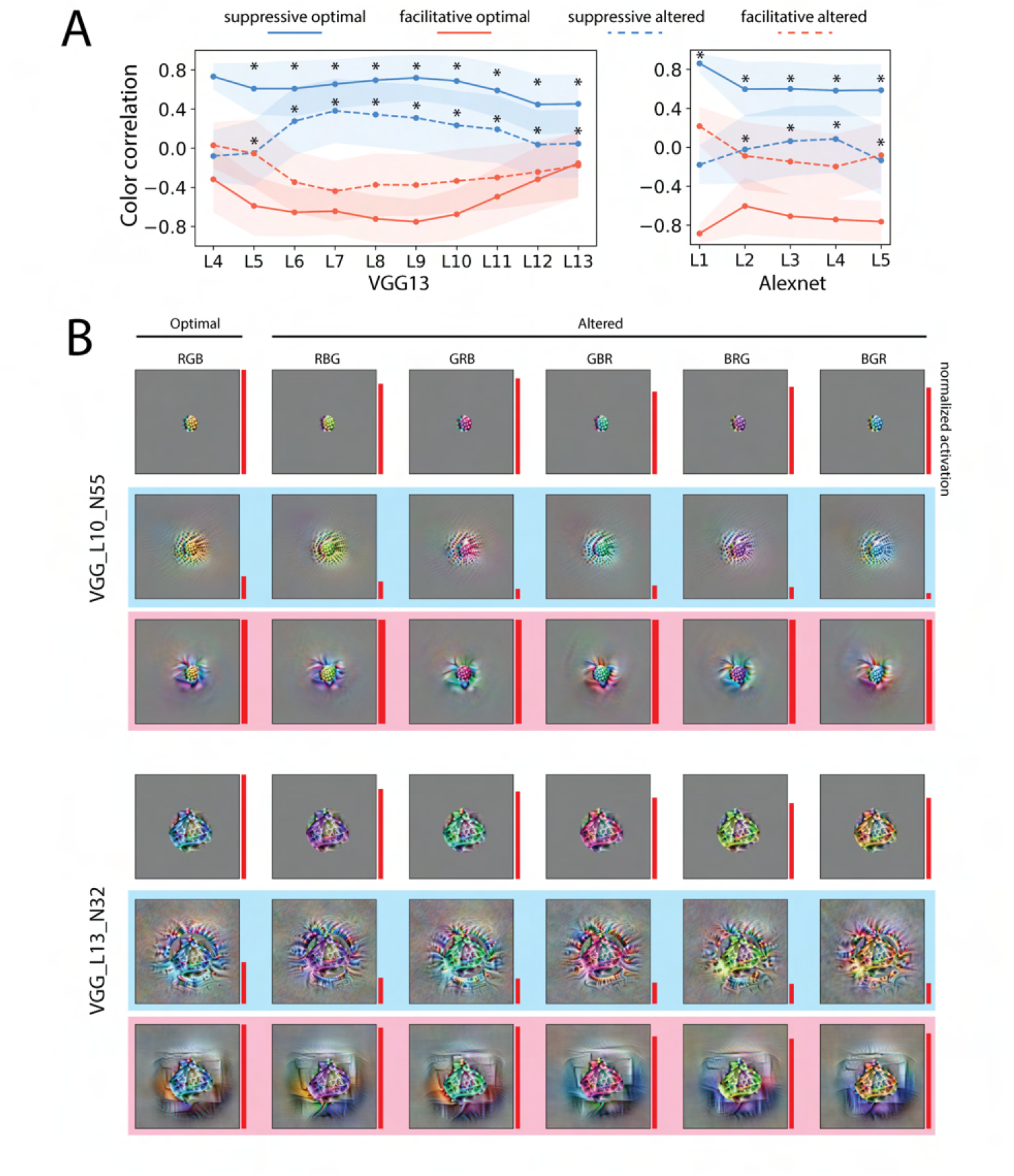
The most suppressive surround can follow the center color change. A. Averaged color correlation of the center the surround in VGG16 (left) and Alexnet (right). Higher values indicate higher color similarity between the center and surround. Four conditions are shown in the plot: the correlation between the optimal center and the most suppressive surround (solid blue); the optimal center and the most facilitative surround (solid red); the altered center and the most suppressive surround (dotted blue); the altered center and the most facilitative surround. The optimal center is defined as the most facilitative center. The altered center is the optimal center with three color channels permuted. The shaded area indicates the standard deviation. B. Two example neurons (VGG L7 N7 and (VGG L10 N55)) showing that the most suppressive surround can match the center color. For each neuron, the first row are the center stimuli; the second row are the center stimuli with the most suppressive surround; the third row are the center stimuli with the most facilitative surround. The first column is the optimal center; other columns are the optimal center with the three color channels permuted. The area of the red bars on the right of each image represents the normalized response (relative to the optimal center response).

We then further tested the idea of homogeneity by exchanging the entire optimal center. Some CNN neurons showed an ability to match the surround to the exchanged center stimuli. Figure 6A shows an example neuron that had such ability. Its optimal center appeared as purple curves; when the center was changed to triangles, yellow curves, and blobs, the most suppressive surround could match the altered center pattern. Figure 6B shows 5 neurons (including the one in Figure 6A) in VGG16 layer 10. The leftmost column shows the optimal center for each neuron. The 5 optimal centers were used for each neuron to derive the most suppressive surround stimuli. By looking at the columns, we see that the most suppressive surround depends on the center stimuli. Furthermore, some neurons could match the surround to the altered center stimuli.

**Fig 6.**
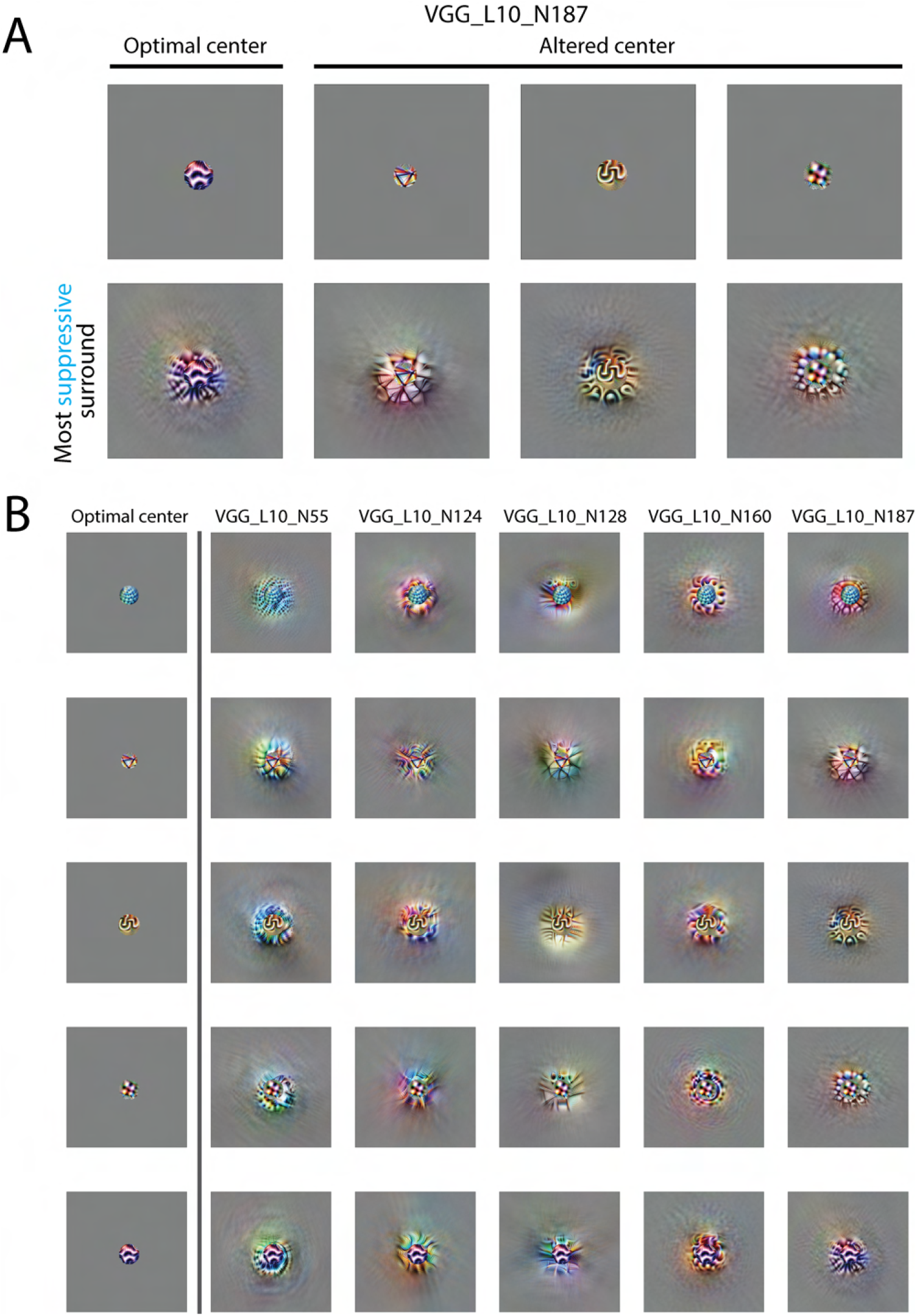
The most suppressive surround depends on the center pattern. A. An example neuron in VGG16 layer 10. The first row is the center stimulus used to optimize the most suppressive surround; the second row is the most suppressive surround using the corresponding center in the first row. The first column is this neuron’s optimal center; the remaining columns are center patterns from the other neurons. B. Visualizations of most suppressive surround of 5 neurons with exchanged centers. The first column shows 5 optimal centers for the 5 selected neurons. Other columns are the most suppressive surround with different centers. The visualizations on the diagonal line used neurons’ own optimal center. The most suppressive surround strongly depends on the center pattern. Some most suppressive surrounds visually matched the center pattern.

Our results suggest that the findings that the most suppressive surround orientation follows the center stimulus in Figure 4 [15, 48] can be generalized to more complex stimuli in the CNN neurons. Such effects with complex stimuli have not been tested in cortical neurons, and therefore provide a testable hypothesis of surround effects in higher visual cortices.

How can the most suppressive surround in the CNN follow the change in the center? This surprising observation in CNNs can be conceptually explained by stacking two layers (Figure 7). Assume that the most suppressive surround of the previous layer is similar to the preferred center, but that it cannot follow a change in the center stimulus (i.e. when the neuron is presented with a non-optimal center stimulus). In the next layer, the most suppressive surround can gain this ability due to the nonlinear activation function after the previous layer. In detail, one center stimulus elicits an activation profile in the previous layer; the most suppressive surround should match this profile to gain the maximum suppression. Thus, the surround matches the center pattern. Otherwise, if the most suppressive surround stays the same as the preferred center rather than the altered center pattern, the excessive suppression to some neurons in the previous layer will not be passed due to rectification of the ReLU activation (Figure 7). A conceptual model with two neurons and a simulation of the visualization experiment (using the same visualization algorithm that was used to generate other figures) are shown in Figure 7. A similar model has been proposed for retinal ganglion cells to explain other surround effects [67]. Though theoretically, two layers can achieve this ability, in practice more layers may be required according to how the assumption is satisfied. This may explain why we only see this ability clearly in later layers in VGG16 but not in early layers or in Alexnet which is shallower.

**Fig 7.**
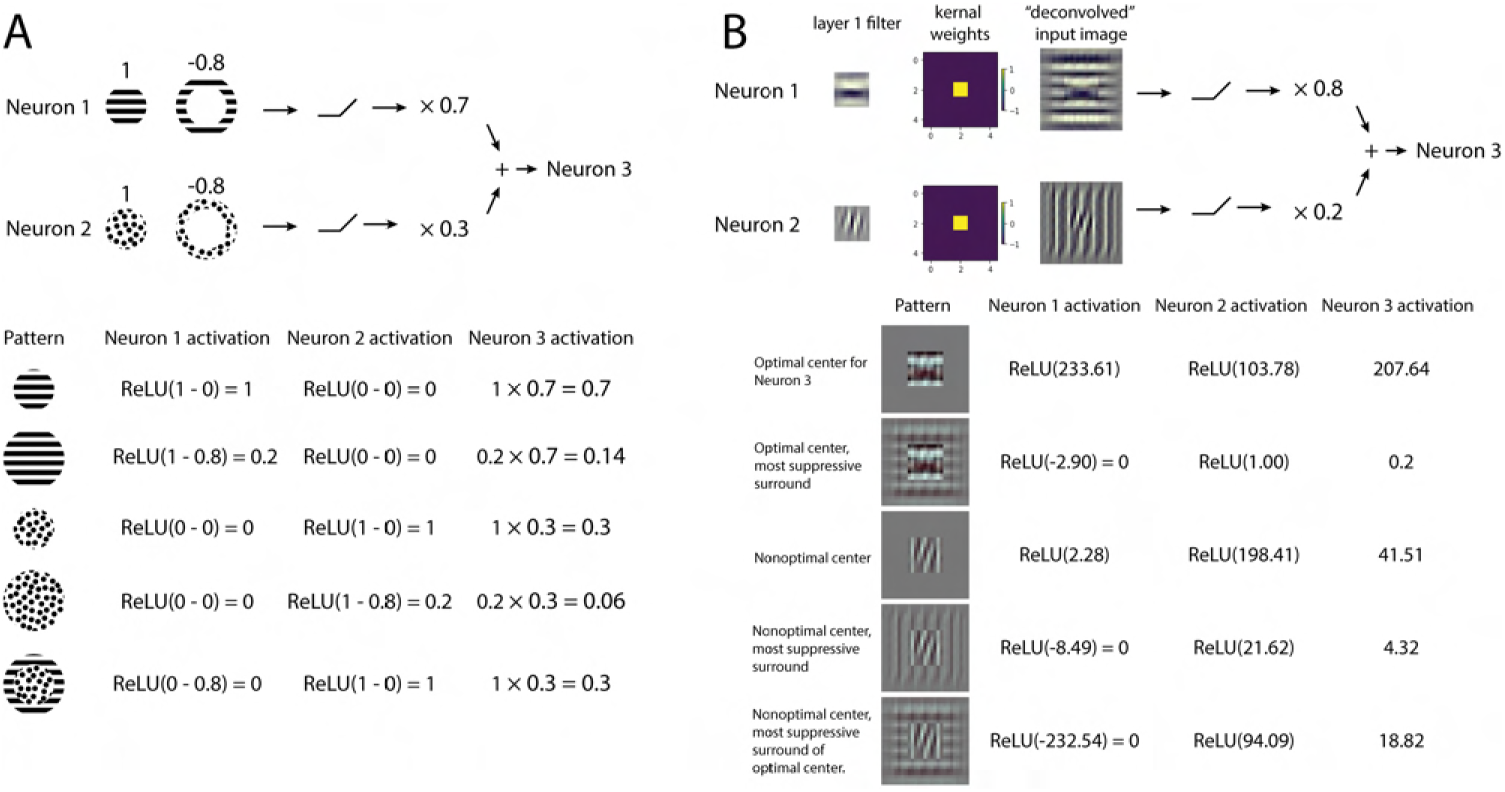
The most suppression by the homogeneous surround can be achieved by two layers with nonlinear activation, as shown in a conceptual model and toy model simulation. A. A conceptual model. Top: The model has two input neurons with center-surround receptive fields, i.e. the optimal pattern in the center elicits a response of 1; the optimal pattern in the surround elicits a response of -0.8. We further assume that the optimal patterns for two neurons are orthogonal, i.e. the optimal pattern of one neuron does not elicit responses of the other neuron. The response of the output neuron, i.e. neuron 3, is the weighted sum of rectified responses of the two neurons. Neuron 1 has a higher weight than neuron 2. Bottom: input patterns that are created from combining optimal patterns of neuron 1 and 2. Activations of three neurons are computed as defined. The maximum surround suppression happens when the surround pattern matches the center. This is due to the nonlinear ReLU blocking the excessive suppression from the unmatched neuron. B. In a toy model simulation, the visualization results show similar behavior to the conceptual model. The two neurons are constructed by first taking two simple orientated filter maps, then convolving with a kernel with center weight 1 and surround weight -1. Equivalent filters for the two neurons are also shown as ”deconvolved” input images. We simulated the visualization experiment in neuron 3. The optimal center for neuron 3 is horizontal and the most suppressive surround matches the optimal center. When the center is replaced by the neuron 2’s optimal pattern, the most suppressive surround looks vertical which is close to the optimal pattern of neuron 2, unlike the most suppressive pattern when the center is fixed to the optimal.

An alternative explanation for the most suppressive surround following the center has been suggested in cavanaugh2002b. If the surround is not entirely isolated and impinges onto the center, then the effect of including a non-optimal center and an optimal (unmatched) surround would be that the surround both excites and suppresses the neural responses; if the additional excitation exceeds the suppression, then this could result in overall less suppression. In contrast, the non-optimal center with the matched non-optimal surround could yield an overall more suppressive balance between the excitation and suppression. However, if this were the complete explanation, it’s not clear why it would be more prevalent in higher layers of the CNN.

### Texture induces less surround suppression than spectrally matched noise

There have been limited studies on surround effects beyond V1. When using grating stimuli, V2 neurons have shown some similar properties to V1 regarding surround effects [68]. Textures that extend beyond the classical receptive field can result in surround suppression in both V2 [31] and V4 [32], an observation that has been referred to as ”de-texturization”. However, other observations in V2 cannot be simply explained by surround suppression based on the center surround similarity, and rather depend on whether the stimulus is naturalistic or noise. In particular, V2 neurons show less surround suppression for naturalistic textures (that include dependencies across space) than for spectrally matched noise [31] (Fig 8D). We therefore asked whether CNN neurons could capture such effects, following a similar design [31]. We synthesized 225 naturalistic texture images from 15 original texture images and their corresponding spectrally matched noise images (see Methods). The optimal textures for each neuron were determined by finding the modulation indexes, i.e. the difference of the responses to the naturalistic and noise images divided by the sum of the two (Fig 8B). The top 5 textures for each neuron were used for the following experiment. For each neuron, we computed the naturalistic and noise diameter tuning curves (Fig 8C, E). We then computed the suppression index (SI), i.e. the difference of the max and min response divided by the max response, for the naturalistic and noise stimuli from the diameter tuning curves.

**Fig 8.**
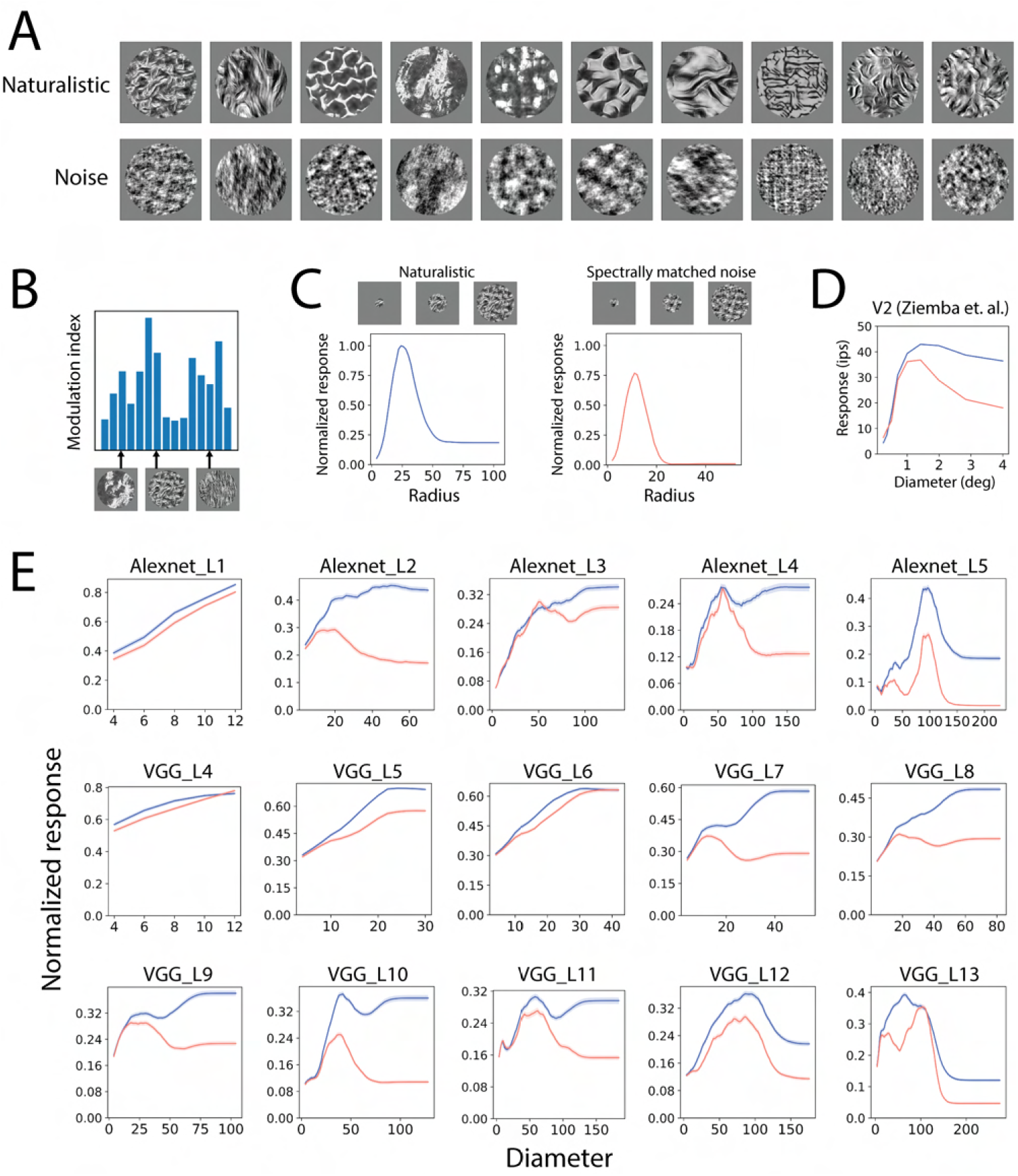
Surround suppression of naturalistic textures and noise. A. Naturalistic textures and spectrally matched noise used in the experiments. Naturalistic textures were synthesized using an algorithm described in Methods. B. Texture tuning of an example CNN neuron. The ”optimal” textures for each CNN neuron was determined by the textures with the highest modulation index (see details in Methods). The ”optimal” textures were then used to study the texture surround effects. C. Texture and noise diameter tuning curves for an example CNN neuron. D. Averaged naturalistic texture and spectrally matched noise diameter tuning curves in monkey V2 (Reproduced from [31]). Noise induces stronger surround suppression. E. Averaged diameter tuning curves in CNN layers. Noise appears to induce stronger surround suppression in most layers.

The CNN neurons exhibited more surround suppression for the naturalistic textures than for the spectrally matched noise, except for early layers of the CNNs (Alexnet layer 1 and VGG16 layer 4). This was observed in both the averaged diameter tuning curves (Fig 7E) and scatter plots (Supplementary Fig 10) of the suppression index. Such effects were consistent with the V2 neurophysiology data (Fig 7D). This result is interesting because it could not be explained by divisive normalization models and is not expected from a model that focuses on the homogeneity of center and surround. We found that it arises spontaneously in the CNN from a stacking of layers. Intuitively, noise images contain no class information, so it makes sense that the network would learn to suppress noise more than textural images.

### Failures of capturing cortical contextual surround effects in CNNs

Though in the above experiments we found some striking commonalities between the CNN neurons and cortical neurons regarding surround effects, we also found failures of the CNNs. One mismatch we found is related to the geometric structure of the surround. Cortical neurons show the largest suppression when the stimulus in the surround is in the location that aligns with the orientation [15] (Fig 9A). We did not find this effect in CNN neurons. Though in some layers responses were significantly different with different surround locations, the effect size was small compared to the biology (Fig 9C). In particular, when the orthogonal surround was used, the trend did not match biology (Fig 9C). We also did not find individual neurons that matched the biological trend (Fig 9B). See Supplementary Figure 11 for the average effects and population distribution of all the layers.

**Fig 9.**
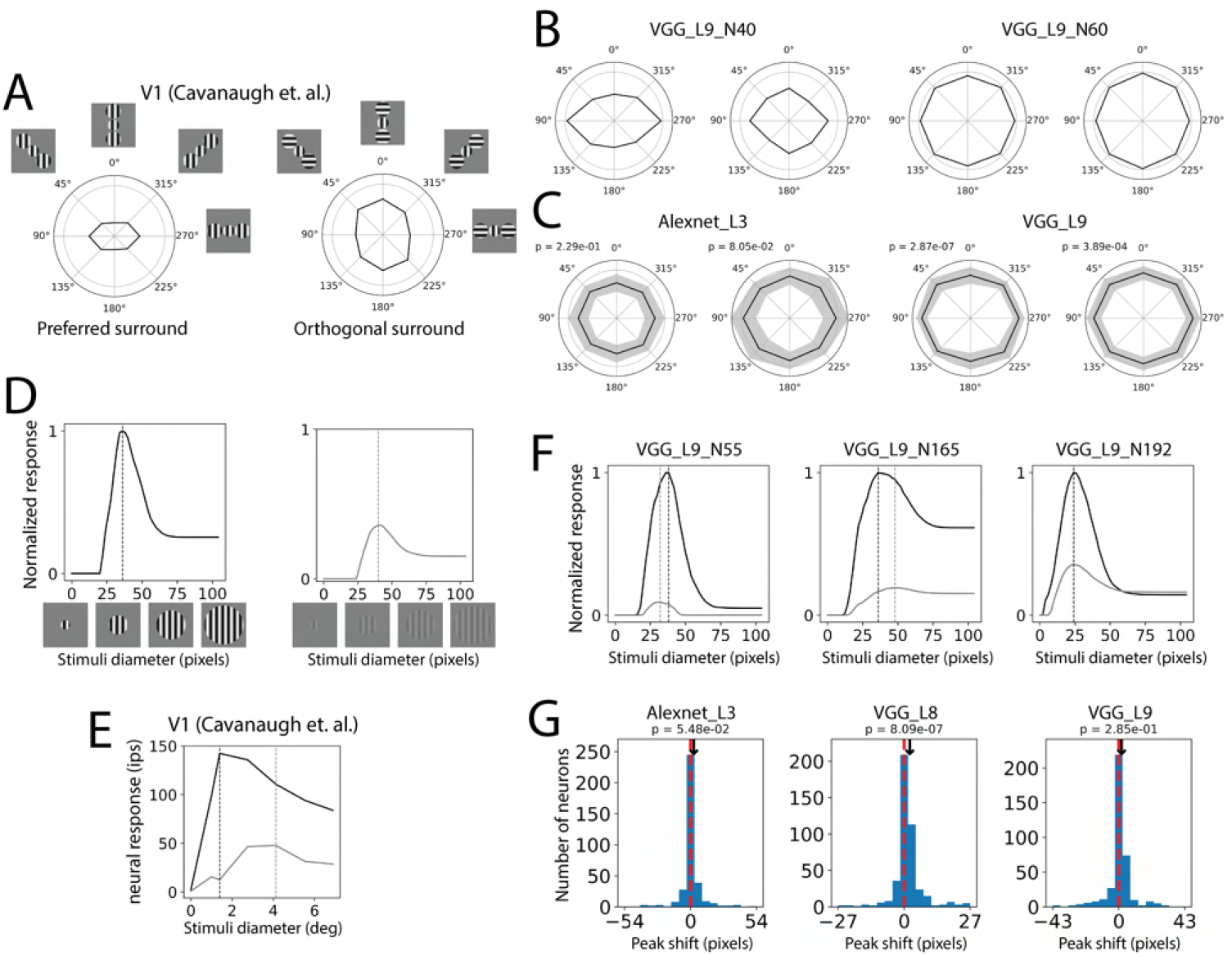
Two mismatches between CNN neurons and cortical neurons. A, C: Geometry effects of the surround suppression. A. Strength of the surround suppression depends on the location of the surround stimuli. Embed images show the stimuli used in this experiment. The center was fixed at the optimal orientation. The surround had two patches at different locations relative to the center stimuli. The surround was either at optimal orientation or orthogonal orientation. Surround patches that align with the center stimuli induce the strongest suppression when it is at optimal orientation and the strongest facilitation when it is at orthogonal orientation. The polar radius represents the normalized response where the gray circle represents 1 (reproduced from [15]). B. Plots of two example CNN neurons. C. Averaged plots of two CNN layers. P values were calculated from one-way repeated measure ANOVA. Though some neurons and layers showed significant modulation effects of surround location, the effect size and shape of the plots did not match the cortical neurons. D, E, F, G: Peak shift of the low contrast diameter tuning curve. D. We computed diameter tuning curves of each neuron with the normal contrast (high contrast, black line) and 17% of the normal contrast (low contrast, gray line). Dotted vertical lines indicate peaks of the diameter tuning curves. E. Two diameter tuning curves of an example V1 neuron (reproduced from [15]). The low contrast peak is shifted rightward. F. Three example CNN neurons with different directions of peak shift. G. Histogram of peak shift in three CNN layers. Peak shift is defined as low contrast peak diameter subtracting high contrast peak diameter. Positive values are more commonly found in cortical neurons. P values were calculated from paired t-test.

Other mismatches we found are related to the contrast. Neurophysiology studies have shown that the grating diameter tuning curve peaks later when the contrast is low [10, 16] (Fig 9 D, E). Though some individual CNN neurons showed this effect, we did not find this consistently in the CNN neurons (Fig 9 F, G, Supplementary Fig 8). Only layer 8 in the VGG16 showed significantly more neurons where the low contrast stimuli shift peak later (Fig 9 G, Supplementary Fig 8). See Supplementary Figure 12 for the averaged curves and peak shift histogram of the layers.

## Discussion

We studied visual contextual surround effects in CNN neurons. First, we simulated a classic visual surround effects experiment in CNN neurons and found that the most suppressive surround grating orientation matches the optimal center orientation (Fig 2). Second, we developed a method to visualize the surround effects in CNN neurons and found that the most suppressive surround is visually similar to the optimal center pattern (Fig 3). The visualization experiments could be thought of as a generalization of the classic grating experiments, but with complex stimuli. We also found that for both grating (Fig 4) and more complex stimuli per the visualization (Fig 5) experiments, the most suppressive surround in deeper layers can still match the center even when the center is non-optimal. The finding for more complex stimuli presents an interesting prediction that could be tested experimentally.

In recent years, optimization-based neuron control techniques have been used in neuroscience experiments to find stimuli that elicit the strongest neural responses and even control the population activation pattern [42–44]. Optimization techniques similar to what we have shown here for the CNNs could be modified to find the most suppressive surround stimuli in neurophysiological studies (since it is extremely difficult to calculate the gradient in the brain, one would need to replace the gradient-based optimizer with a gradient-free optimizer such as Covariance Matrix Adaptation). Such findings could reveal new surround effects across the visual hierarchy, and help elucidate to what extent paradigms of homogeneity play a role in surround suppression in higher visual areas.

How can a generic feedforward CNN capture a signature of surround suppression? Since surround effects are thought to arise from nonlinear computations such as divisive normalization and recurrent and feedback connections, feedforward CNNs without those connections are not supposed to capture such effects. However, CNNs can potentially capture surround effects by stacking layers. From a statistical perspective, this could relate to computational studies showing that in deeper layers of CNNs, the activations of neighboring neurons are less statistically dependent, thereby achieving some of the statistical properties that have been attributed to divisive normalization [69]. Surround suppression in CNNs may also be achieved by subtractive suppression from the surround, due to a combination of weighted outputs from the previous layer (i.e., more negative weights on average in the surround). We did find that in early layers, the center-surround receptive field may arise from the convolution kernel weights having a center-surround structure (Supplementary Figure 15).

Is the surround suppression we observe in standard feedforward architectures subtractive or divisive? Some studies have shown that surround effects in neurophysiology are better fit with a divisive than subtractive model [16, 70, 71]. Contrast experiments have been used to distinguish if the surround effects are divisive and/or subtractive. We simulated an experiment with such a design (Supplementary Fig 13). The results showed a preference for the subtractive model over the divisive model especially in early layers (Supplementary Fig 14). Though generic CNNs do not contain division explicitly, our results rejected that the surround effects are purely subtractive. Elucidating the abstract mathematical form of surround effects in CNNs will require future studies.

Studies in V2 have shown that there are additional factors that influence surround suppression, namely whether the stimulus itself is naturalistic or noise. In particular, there is less surround suppression for extended natural textures than for spectrally matched noise [31] (Fig 8), suggesting that the brain suppresses noise stimuli more than naturalistic stimuli. This behavior is conceptually reasonable, since noise contains less information about the image classification and so is better to be suppressed. This result cannot be explained by divisive normalization models based on center surround homogeneity, but is interestingly captured in CNNs.

In addition to the success cases, we also found important mismatches between CNN neurons and cortical neurons. First, we point out that in the generic CNN model, there is not a clear separation of center and surround regions as in the visual cortex (see Methods). The surround alone in our simulations sometimes elicited a response unlike the convention in neurophysiology (see Figure 2). In terms of the simulations, surround suppression was less dependent on the geometric location of the surround stimuli in CNN neurons than in the neurophysiology (Fig 9, Supplementary Fig 11). Our notion of homogeneity in this study is indeed more limited than image statistics models that infer the statistical dependencies between the center and surround stimuli, and therefore can capture such geometric effects [21]. This indicates that CNNs do not capture natural scene statistics pertaining to the geometry, or that this type of geometric dependency is not required for the image classification task which the neural network was trained on. This suggests that explicitly incorporating such scene statistics in deep neural networks via divisive normalization [26, 50] may improve the results. We also found that for contrast changes, CNNs behave differently from cortical neurons. In the brain, the grating diameter tuning curve peaks later when the contrast is low [10, 16, 27], suggesting, for instance, that there is broader facilitation rather than suppression of lateral inputs when the inputs are weak [29]. Though this emerged for some CNN neurons, it did not reach statistical significance in most layers of the CNN (Fig 9, Supplementary Fig 12). This may be due to the activation function used in CNNs (in our case, ReLU), rather than saturating functions that could emerge from divisive normalization. Such contrast phenomena have been explained by image statistics models [21, 24], and again suggest routes for improving the results of CNNs in future work. Another aspect that the model did not capture is the greater suppression for gratings than for texture and noise stimuli [31], which may again require a mechanism for contrast normalization.

The connection between surround effects and neural coding has been studied in theoretical works [12, 13, 20, 22, 24] example, surround effects may be explained from an efficient coding perspective via a single layer divisive normalization model that is derived from reducing high-order dependencies [21, 26, 72]. Empirically, in deep neural networks, we have found that spatially neighboring artificial neurons with the same feature selectivity have statistical dependencies (and higher mutual information) in lower layers, but less dependencies (and lower mutual information) in higher layers [50, 69]. This has led us to hypothesize that deep networks may reduce statistical dependencies by stacking layers. More recently, a formal connection between deep learning and (a different form of) efficient coding has also been suggested in CNNs: networks learn more common features faster and are more sensitive to them [73]. However, this has not been linked to spatial surround statistics or surround effects. In future work, we would like to test the hypothesis that the surround effects we found in CNNs arise from high-order spatial dependencies in natural image statistics and can be captured by gradient descent learning. However, the current theory is limited to the analysis of linear networks with first-order statistics. Future works are needed to extend those theories to non-linear networks with high-order statistics to test our hypothesis.

Surround suppression has been considered to have a number of beneficial roles in neural computation, for example, reducing coding redundancy and highlighting salient features [12, 14, 19–22, 24, 26, 28, 72]. Some studies in machine learning noticed the lack of more sophisticated forms of brain-like divisive normalization in generic feedforward CNNs, and tried to integrate them into the network [49–53]. These studies found that incorporating divisive normalization in CNNs improves image classification in some limited cases, such as when the network is more shallow [51], when the dataset requires strong center-surround separation [51], or when the divisive normalization is combined with batch normalization [52]. The correspondence we found between generic CNNs and the brain regarding center surround similarity may explain why including divisive normalization explicitly in CNNs has only limited improvement in classification, especially when the networks are deep. However, some of the mismatches also suggest a need for exploration of such deep learning architectures that explicitly incorporate contextual information. This is in line with other studies showing that complex perceptual uncrowding phenomena are not explained by generic CNNs and require a mechanism for grouping and segmentation [74]. Studies of contour integration have also incorporated functional columns and lateral connections [54, 75]. Biologically inspired computations that efficiently capture surround effects may help design artificial neural networks that are shallow and more efficient.

Our findings overall demonstrate that standard feedforward architectures exhibit surround suppression based on the similarity between center and surround stimuli, suggesting that such architectures can capture and generalize an important characteristic of surround effects in cortical neurons. The visualization approach could be adopted to study surround effects in high visual areas. The mismatches we found may inspire future studies of contextual effects in deep neural networks with more sophisticated circuitry, including the role of divisive normalization [49–53, 76], recurrent connections and feedback [77–82] in hierarchical architectures.

## Methods

### CNN models

We trained an Alexnet-style and a VGG16-style network on the Imagenet dataset mostly following the original papers respectively [55, 56]. Model files are available in the online repository. Our results are not altered qualitatively when using other publicly available CNN instances. There are several changes we made to the original model.

These changes are prevalent and have become almost new standards. We replaced local response normalization in Alexnet with batch normalization, and step decay with cosine decay for the learning rate scheduling. For training Alexnet, we trained on one GPU rather than two as in the original study. We used the data augmentation process described in [55] for training the CNNs. We applied the standard Xavier uniform method to initialize weights in the convolutional and dense layers. The architectures are shown in table 1. Since the dense layers do not have spatial feature maps that are crucial for determining surround effects, only convolutional layers are analyzed in this study.

**Table 1.** CNN architectures used in this study. The input size of both networks is 224”224”3. Conv2D represents 2D convolutioanl layer. Three following numbers denotes the kernel size, stride size and channel numbers. BN represents batch normalization. MaxPooling represents 2D max pooling layer. The following numbers denotes the pool size and stride size. Dropout represent dropout layer. The following number denotes dropout rate.

|  |  |
| --- | --- |
| Alexnet | VGG16 |
| Conv2D_11_4_96-BN-ReLU (L1) | Conv2D_3_1_64-ReLU (L1) |
| MaxPooling_2_2 | Conv2D_3_1_64-ReLU (L2) |
| Conv2D_5_1_256-BN-ReLU (L2) | MaxPooling_2_2 |
| MaxPooling_2_2 | Conv2D_3_1_128-ReLU (L3) |
| Conv_2D_3_1_384-BN-ReLU (L3) | Conv2D_3_1_128-ReLU (L4) |
| Conv_2D_3_1_384-BN-ReLU (L4) | MaxPooling_2_2 |
| Conv_2D_3_1_256-BN-ReLU (L5) | Conv2D_3_1_256-ReLU (L5) |
| MaxPooling_2_2 | Conv2D_3_1_256-ReLU (L6) |
| Dense_4096-BN-ReLU | Conv2D_3_1_256-ReLU (L7) |
| Dropout_0.4 | MaxPooling_2_2 |
| Dense_4096-BN-ReLU | Conv2D_3_1_512-ReLU (L8) |
| Dropout_0.4 | Conv2D_3_1_512-ReLU (L9) |
| Dense_1000-BN-ReLU | Conv2D_3_1_512-ReLU (L10) |
| Dropout_0.4 | MaxPooling_2_2 |
| Dense_1000-BN-Softmax | Conv2D_3_1_512-ReLU (L11) |
|  | Conv2D_3_1_512-ReLU (L12) |
|  | Conv2D_3_1_512-ReLU (L13) |
|  | MaxPooling_2_2 |
|  | Dense_4096-ReLU |
|  | Dropout_0.5 |
|  | Dense_4096-ReLU |
|  | Dropout_0.5 |
|  | Dense_1000-Softmax |

Since the feature maps of all the convolutional layers in our study have an even number of neurons in height and width, the center neurons we selected are actually a half unit away from the image center. And these half unit displacements in the feature maps correspond to different pixel numbers when tracing back to the input image. In this study, we always put stimuli at the true center of each neuron. That means for each layer, the displacement of the stimuli from the image center was adjusted based on the shape of the feature maps. The displacement in pixels is calculated by the formula:

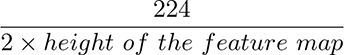

For each CNN neuron, we can derive a theoretical receptive field by tracing the feedforward computations [83]. Note that this theoretical receptive field is different from the classical receptive field in the neurophysiology literature. Stimuli beyond the theoretical receptive field are guaranteed to have no effect on the neuron’s responses. We followed the method in [83] to compute the theoretical receptive field for each CNN layers. We denote, in layer i, the receptive field size *r_i_*, the kernel size *k_i_*, and the stride size *s_i_*. In order to calculate *r* recursively, we need another attribute, jump size *j_i_* in previous layers which is the distance in pixels between two adjacent units. For the input layer, *j*_0_ = 1, and *r*_0_ = 1. *j* can be recursively computed, *j_i_* = *j_i−_*_1_ *∗ s_i−_*_1_. Then *r* can also be recursively computed, *r_i_* = *r_i−_*_1_ + (*k_i−_*_1_ *−* 1) *∗ j_i−_*_1_. The above two recurrence equations can be solved, 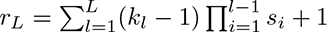. The values are shown in Supplementary Figure 1. The code can be found in the online repository.

### Finding optimal grating stimuli for each CNN neuron

A key component of our simulations was to define the diameter that separates the center and surround; in other words, an analogy to the classical receptive field in neurophysiology settings. We tried to mimic the neurophysiology experiments that define these diameters as much as possible [15] (Fig 1A). First, we characterized neurons by their optimal grating orientation and spatial frequency by grid searching. We used 3 different stimuli sizes: 30%, 50%, and 70% of the theoretical receptive field described in the previous section; 24 spatial periods (the reciprocal of spatial frequency) from 4 pixels to 50 pixels; and 12 orientations from 0° to 180°. Each response value of these stimuli is the average of 8 different phases mimicking the drifting effects in some experiments [15]. We defined the optimal grating orientation and spatial frequency of each neuron by the stimulus that maximally activates it regardless of the stimulus size. Then, these optimal parameters were used to get the diameter tuning curve of each neuron. The grating summation field was defined by the smallest diameter that elicits at least 95% of the maximum response [15]. The grating summation field was used as the border of the center and surround in our experiments.

### In-silico simulations

We did in-silico simulations on CNN neurons following the experimental neurophysiology paradigms as much as possible. For each neural feature map in each layer, we only selected the center neuron (see the section ”CNN models”) to do the simulation. The method we used to get optimal grating parameters and define the center region are described in the previous sections.

We did not include all the neurons in the analysis due to either an unsuitable center and surround ratio or lack of response variation in the orientation tuning curve. In detail, in Fig 2, 4, and 9, if a neuron’s grating summation field is smaller than 30% or larger than 70% of the theoretical receptive field, or the center orientation turning or surround suppression curve has less than 0.001 variation, we excluded it in the analysis. This process ensures the selected neurons have biologically plausible response profiles (i.e. excluding silent neurons) and reasonably large center and surround extents to do the simulations. The included neuron numbers are shown in 2. In early layers, due to the small receptive field size, only few neurons were included. Beyond Alexnet layer 2 and VGG16 layer 5, about half the neurons were included. The neural responses were normalized by dividing the optimal center grating stimuli responses for each neuron.

The center and surround border has been described in the previous sections. In more detail, for our simulations, we chose the center diameter according to the grating summation field; the inner diameter of the surround as the grating summation field plus 4 pixels; and the outer diameter of the surround according to the theoretical receptive field size (Fig 2, 4, 9).

Some neurophysiology studies use the diameter tuning curve for the annular stimuli to determine the inner diameter of the surround stimuli [15, 16]. We did not follow this; if we chose the surround extent according to [15, 16] as a 95% reduction in the diameter tuning of the annular stimuli responses, many CNN neurons would have a small surround region. As we noted earlier, in the generic CNN, the center and surround are not entirely separable, and the surround in our simulations elicited a weak response (Fig 2). Instead, we set the inner diameter of the surround as the grating summation field plus a fixed value (4 pixels).

### Feature visualization

One can visualize the features in CNNs by finding the optimal inputs that lead to the maximum activations [45–47]. This optimization is done by using the gradient of an activation target regarding the parameterized inputs. In our case, the optimization targets are the responses of each center CNN neuron before the ReLU activation layer. We chose to optimize responses before ReLU to let the gradient flow better to the input; otherwise, the flat zero part of the activation can cause 0 gradient. If no regularization is applied to the optimization, the visualization is usually biased to high frequency and visually unrecognizable noise. We therefore used two kinds of regularizations: naturalistic power spectrum prior and jitter. In detail, we parameterized the input images into the frequency domain, then used the well-known 1/f power law to rescale the frequency components. For the jitter, images in the spatial domain were randomly shifted in both axes with a maximum value of 8 pixels. These two regularizations can help the visualizations appear more natural. We adapted some code from the python package ”lucid” to do the visualization. Our code is available in the online repository.

The innovation of our visualization method is to visualize the surround effects in two steps: first find the most facilitative center image; then find the most suppressive or facilitative surround image.

In detail, to find the most facilitative center image, the center image parameters were used to construct an image; the surround region (see the previous section for the definition) of the image was replaced by gray; then the resulting image was passed to the CNN. We computed the gradient of a center neuron’s response with respect to the center image parameters. The gradient was used to optimize the center image parameters by the Adam optimizer. This optimization step was repeated for 500 iterations for each neuron.

To find the most suppressive or facilitative surround image, the surround image parameters were used to construct an image; then the center region of this image was replaced by the most facilitative center image described in the previous paragraph; then the optimization procedure for the surround image parameters was the same as the procedure for the center.

We used the Adam optimizer for all the layers in both networks. We found that the learning rate that can generate visualizations with vivid color and clear patterns varies across layers. In Alexnet, the first three layers are 0.001; the latter two layers are 0.005. In VGG16, layers 1 to 5 are 0.0005; layers 6 to 9 are 0.001; layers 10 to 11 are 0.0025, layers 12 to 13 are 0.005.

### Naturalistic texture synthesis

In the naturalistic and spectrally matched noise simulation, we synthesized 225 naturalistic texture images from 15 original texture images and their corresponding spectrally matched noise images [84, 85]. We found the optimal textures for each neuron via the modulation indexes, i.e. difference divided by the sum of responses to the naturalistic and noise images (Fig 8B). We used the top 5 textures for each neuron in the simulations. If there were not 5 textures that could elicit non-zero responses, we only used the textures that could elicit non-zero responses. If no texture could elicit non-zero responses, we dropped the neuron in the analysis. The neural responses were first averaged per texture family for naturalistic or spectrally matched noise respectively. For each neuron, the normalization factor was the maximum of all response values. The responses of a texture family, both naturalistic and spectrally matched noise, were divided by the normalization factor. Then the diameter tuning curves were averaged across neurons to get the averaged diameter tuning curves in (Fig 8E).

The naturalistic and spectrally matched noise images used in this study are generated according to [86]. We used a texture synthesis algorithm that has been applied in many neurophysiology experiments [31, 84, 85, 87]. The algorithm takes an example image as input and a random seed, and iteratively modifies a noise image to match a set of defined image statistics of the example images. The source images we used are from previous neurophysiology studies [85, 87] and include 15 grayscale images with 320 x 320 resolution. We synthesized 15 naturalistic images with different random seeds for each source image. The spectrally matched noise images were generated by replacing the phase of the naturalistic images in the Fourier domain with the phase of Gaussian white noise images.

## Supporting information

**S1 Appendix. Online repository** CNN model files, code, and the full set of the visualization can be found in the online repository: https://gin.g-node.org/xupan/CNN_surround_effects_visualization

---

**S1 Table.** Number of selected neurons. Silent neurons and neurons with too large or small center were excluded from the orientation suppression simulations. But the visualization experiments only excluded neurons with too large or small center and included silent neurons. Silent neurons were defined whose center orientation tuning curve has less than 0.001 variances. Center that is larger than 70% of the theoretical receptive field size was considered too large; smaller than 30% of the theoretical receptive field size was considered too small.

| Layer | Total | Selected | Silent | Center too large | Center too small |
| --- | --- | --- | --- | --- | --- |
| A1 | 96 | 5 | 0 | 78 | 13 |
| A2 | 256 | 123 | 0 | 95 | 38 |
| A3 | 384 | 232 | 5 | 60 | 89 |
| A4 | 384 | 199 | 15 | 23 | 154 |
| A5 | 256 | 82 | 31 | 9 | 142 |
| V4 | 128 | 2 | 0 | 120 | 6 |
| V5 | 256 | 85 | 0 | 163 | 8 |
| V6 | 256 | 127 | 0 | 113 | 16 |
| V7 | 256 | 132 | 0 | 53 | 37 |
| V8 | 512 | 271 | 0 | 101 | 91 |
| V9 | 512 | 250 | 0 | 76 | 186 |
| V10 | 512 | 211 | 3 | 62 | 238 |
| V11 | 512 | 255 | 1 | 55 | 202 |
| V12 | 512 | 254 | 4 | 52 | 203 |
| V13 | 512 | 297 | 3 | 29 | 183 |

**S1 Fig.**
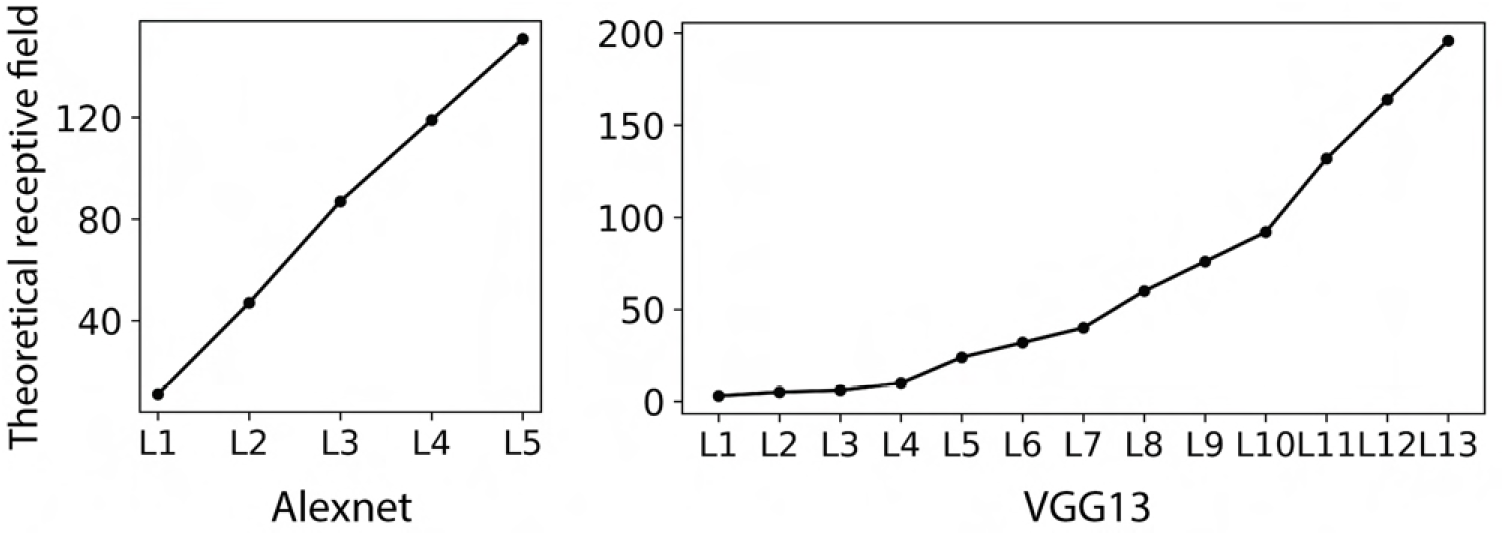
Theoretical receptive field size in VGG16 and Alexnet. See Methods for more details.

**S2 Fig.**
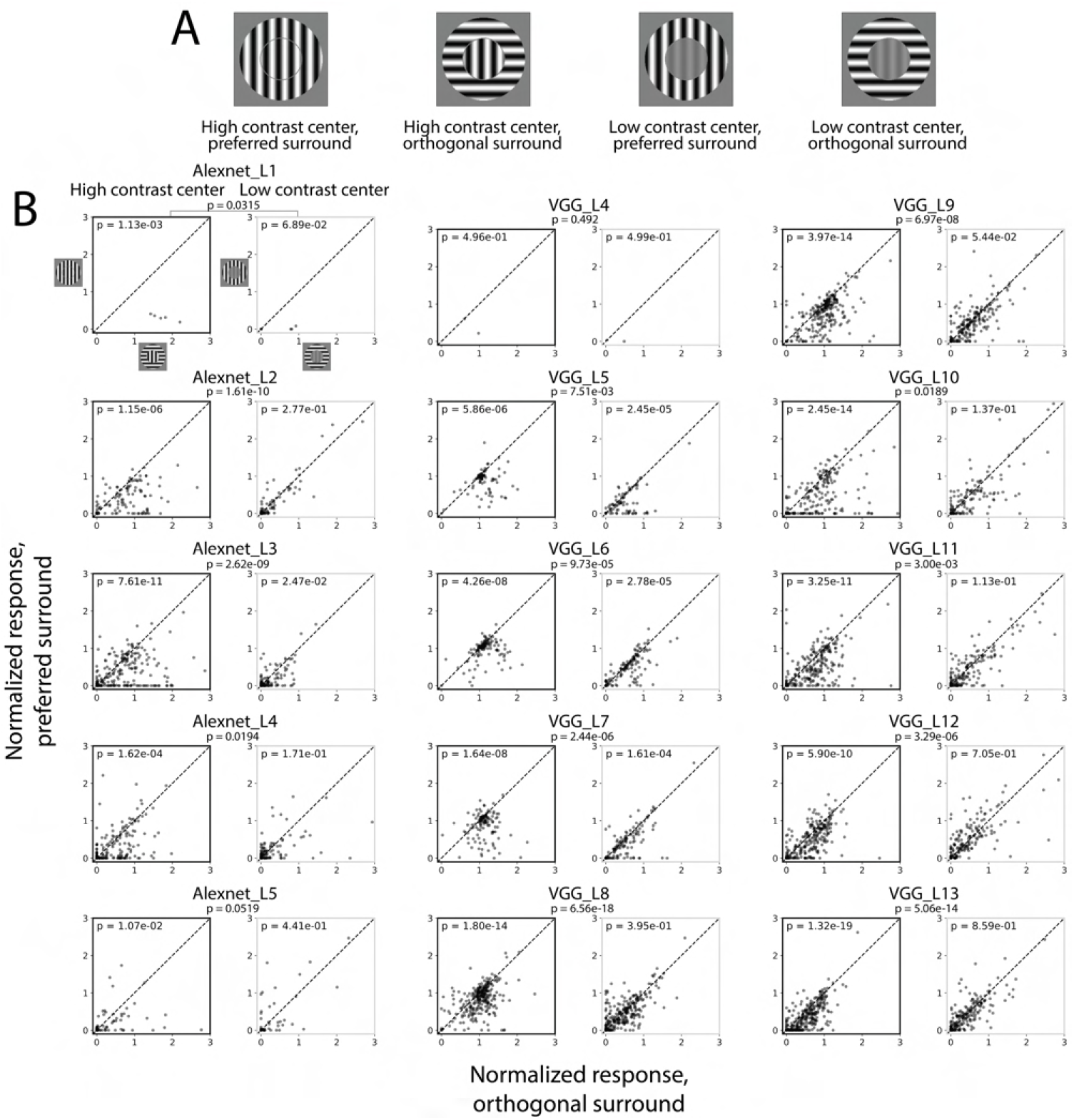
The optimal surround stimulus orientation is more suppressive when the center contrast is high. A. Stimuli used in the experiments. The surround is either at the optimal orientation (preferred surround) or orthogonal to the optimal orientation (orthogonal surround). The center contrast is either at the regular pixel range (high contrast) or 17% of the regular range (low contrast). B. Scatter plots of the responses for preferred surround versus orthogonal surround. Plots with black frames used high contrast center; plots with gray frames used low contrast center. P values inside the plots were calculated from the paired sample t-test (preferred surround versus orthogonal surround). Points below the diagonal lines indicate more suppression when the surround is at the optimal orientation than when it is at the orthogonal orientation. This effect is more pronounced when the center contrast is high. P values outside the plots were calculated from the paired sample t-test of the difference between the preferred surround and orthogonal surround in high contrast and low contrast conditions.

**S3 Fig.**
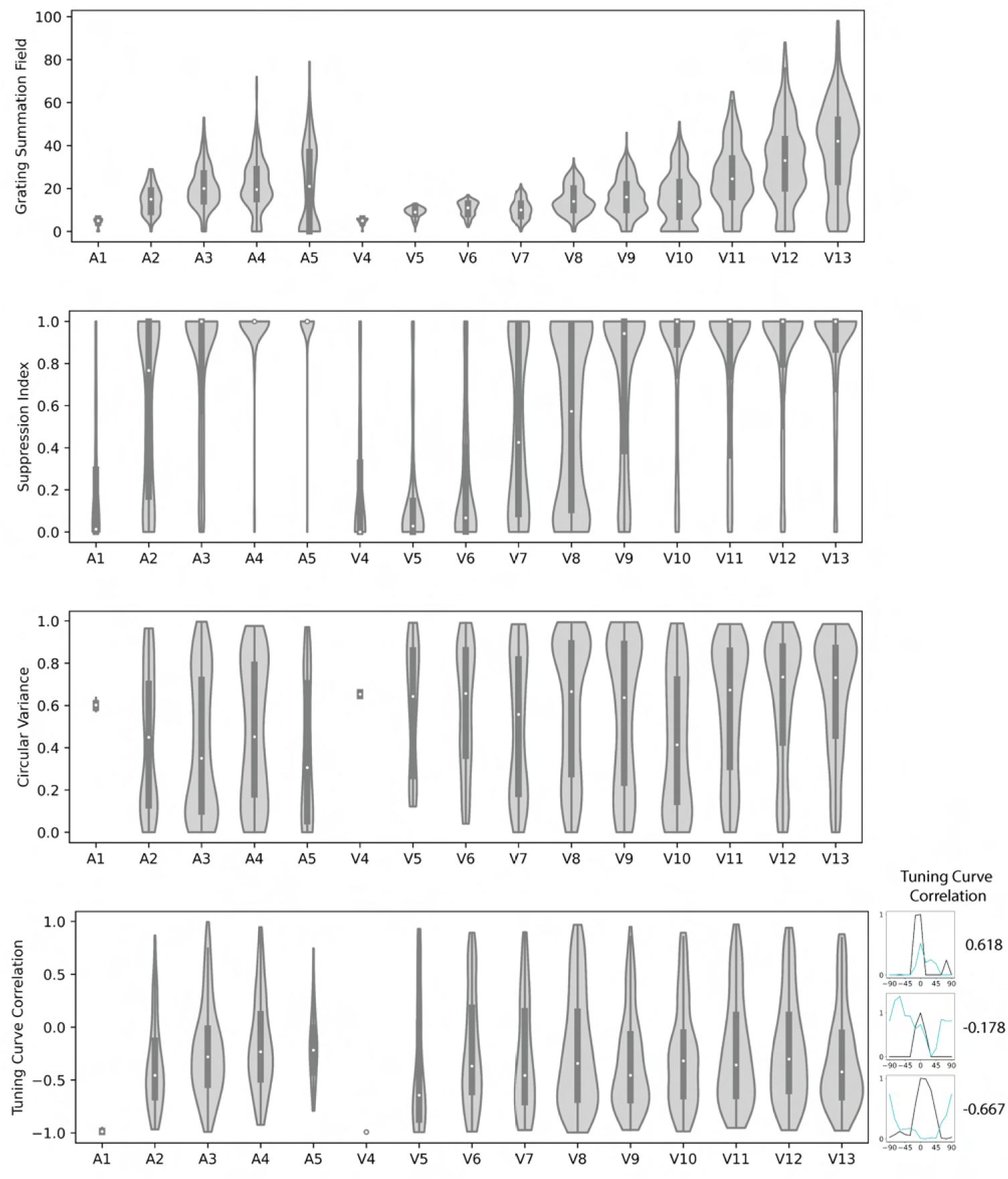
Violin plots of the population distribution of single neuron properties in CNNs. Grating summation field is defined by the smallest diameter that elicits at least 95% of the maximum response. Suppression Index is a metric of surround suppression with grating stimuli which is defined as 1 - (suppressed response / peak response). Circular variance is a metric that characterizes orientation selectivity and is defined in [65]. Small values indicate high orientation selectivity; large values indicate low orientation selectivity. Tuning curve correlation is the Pearson correlation coefficient between the center orientation tuning curve and the surround suppression tuning curve. Correlation coefficients are all negative on average in all layers, which indicates the most suppressive surround orientation matches the optimal center orientation. Three example neurons with different turning curve correlations are shown on the right.

**S4 Fig.**
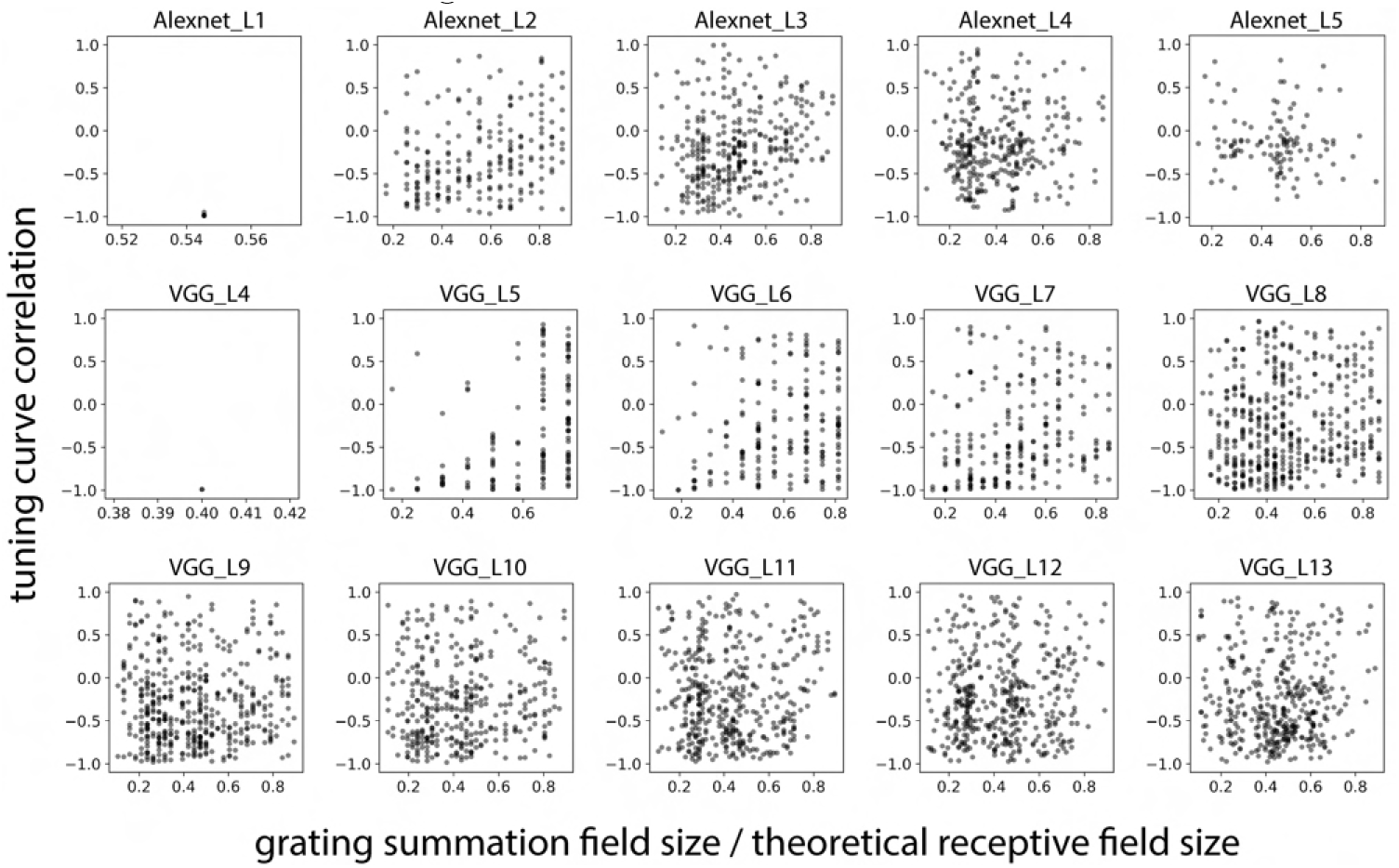
Relationship between tuning curve correlation and the relative size of the center to the surround, i.e. grating summation field divided by theoretical receptive field. In the main experiments, we focused our analysis on the neurons with sufficiently large center and surround sizes. In some layers, especially middle layers, neurons with negative tuning curve correlation are concentrated at a center-surround ratio of 0.2 to 0.5.

**S5 Fig.**
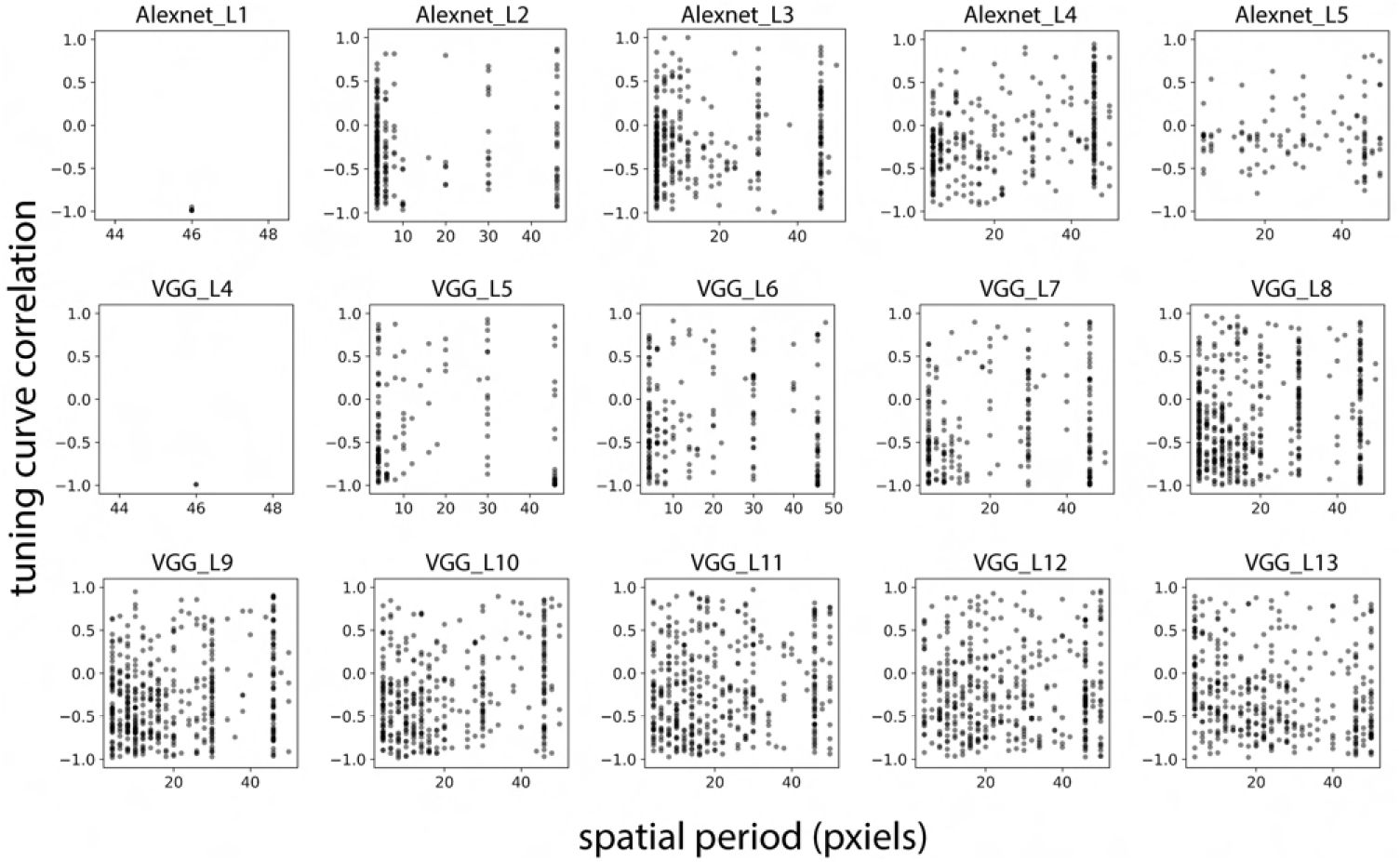
Relationship between tuning curve correlation and the optimal spatial period. We found the distributions are multi-modal. Many neurons with negative tuning curve correlation are concentrated below 30-pixel spatial period. There are some neurons that have large spatial periods, e.g. 50 pixels. Those neurons are likely to be tuned to large color patches or complex patches beyond simple gratings.

**S6 Fig.**
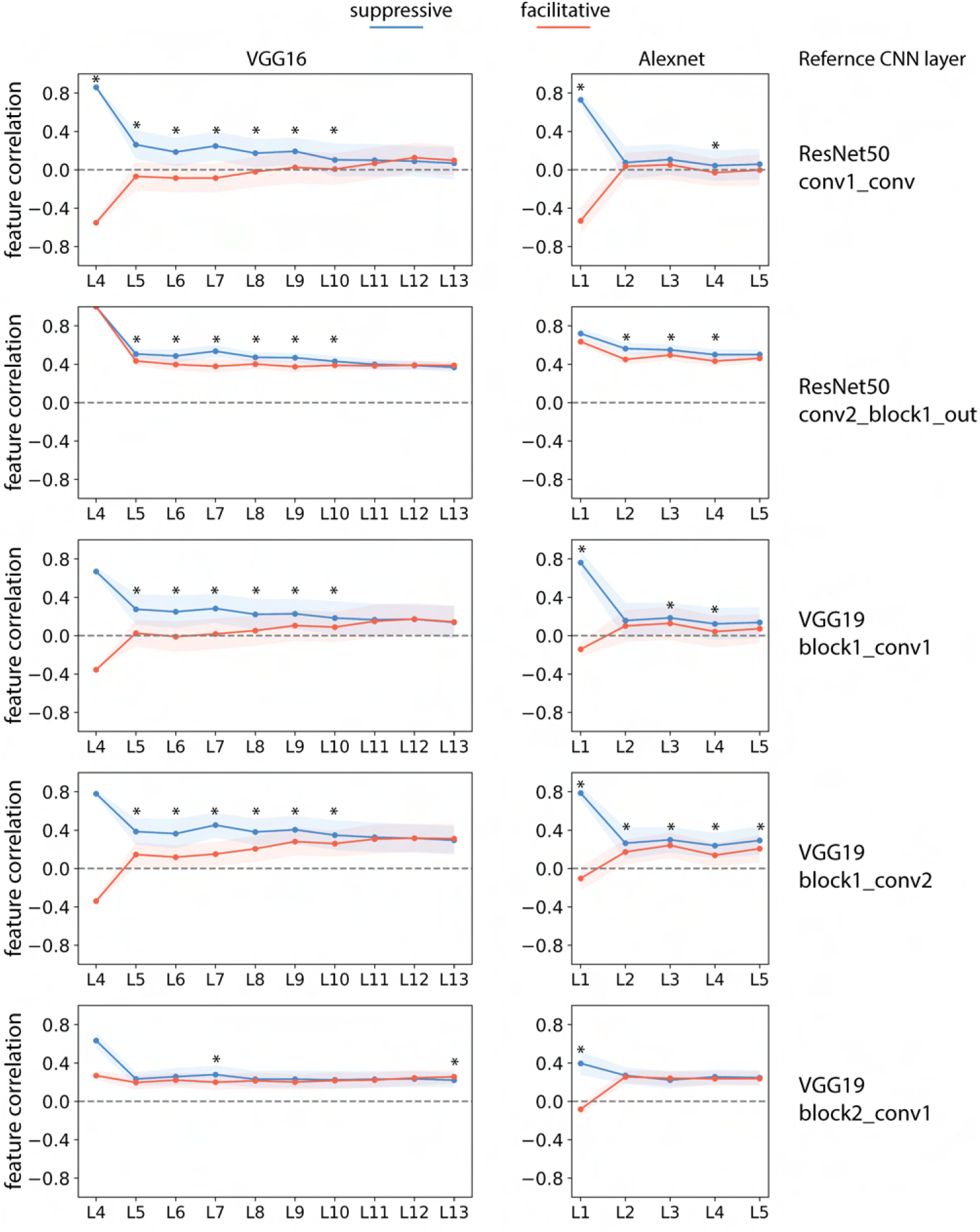
Feature correlation of the center and surround with different reference feature maps. Feature correlation of the center and surround depends on the choice of the reference feature maps. Though the absolute value varies with the feature map choice, they share similar trends that the most suppressive surround has higher feature correlations than the most facilitative surround in early/middle layers in VGG16. Note that a proper choice of feature map should not be a layer that is too deep and has large receptive fields, in which case the surround feature also ”sees” the center. For details on how to compute feature correlation, see Figure 3 caption. The shaded area indicates standard deviation. Asterisks indicate p value smaller than 0.05 in paired t-test.

**S7 Fig.**
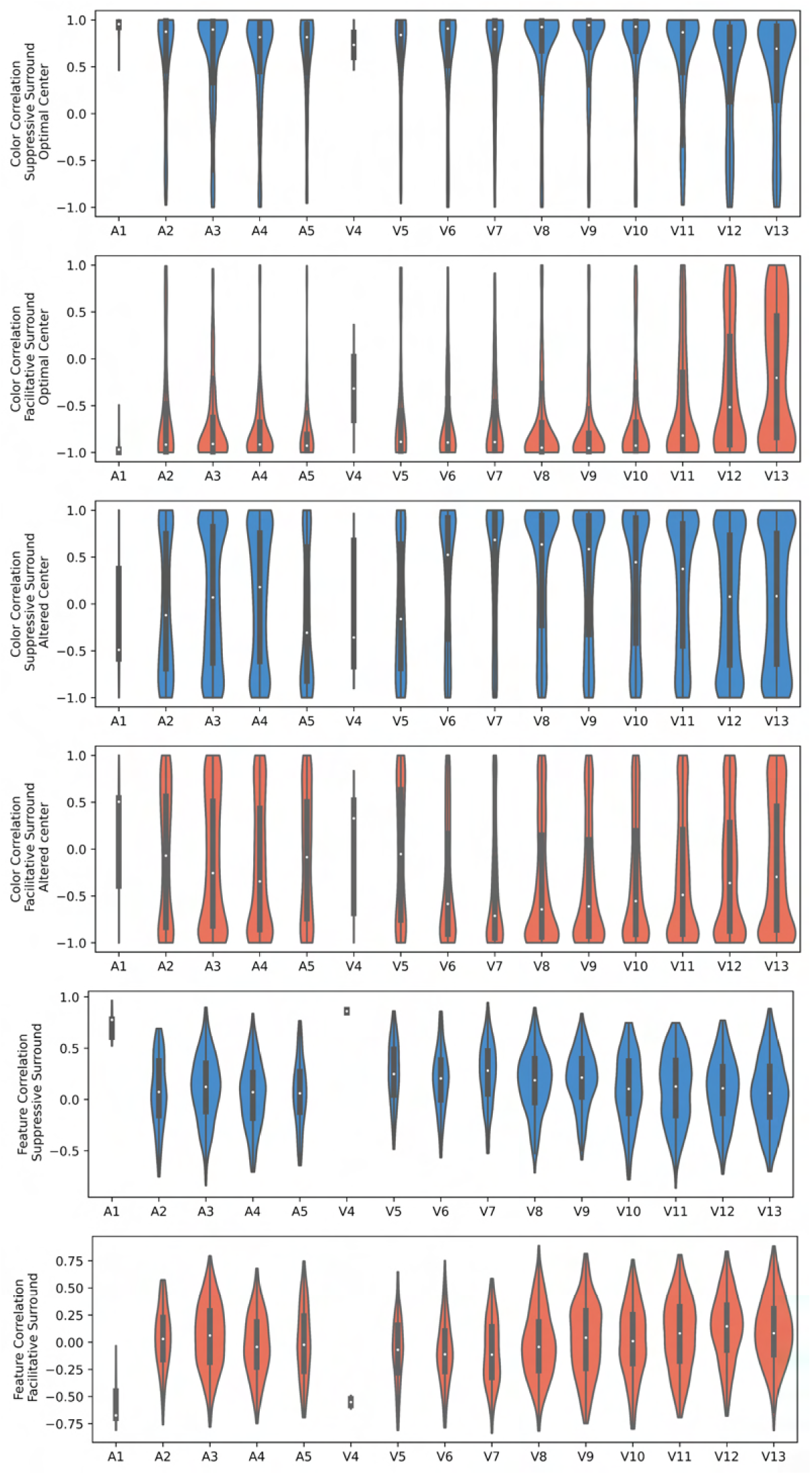
Violin plots of color correlation and feature correlation. Same measure as in Figure 3 and Figure 5 but plotted in violin style to show population distribution.

**S8 Fig.**
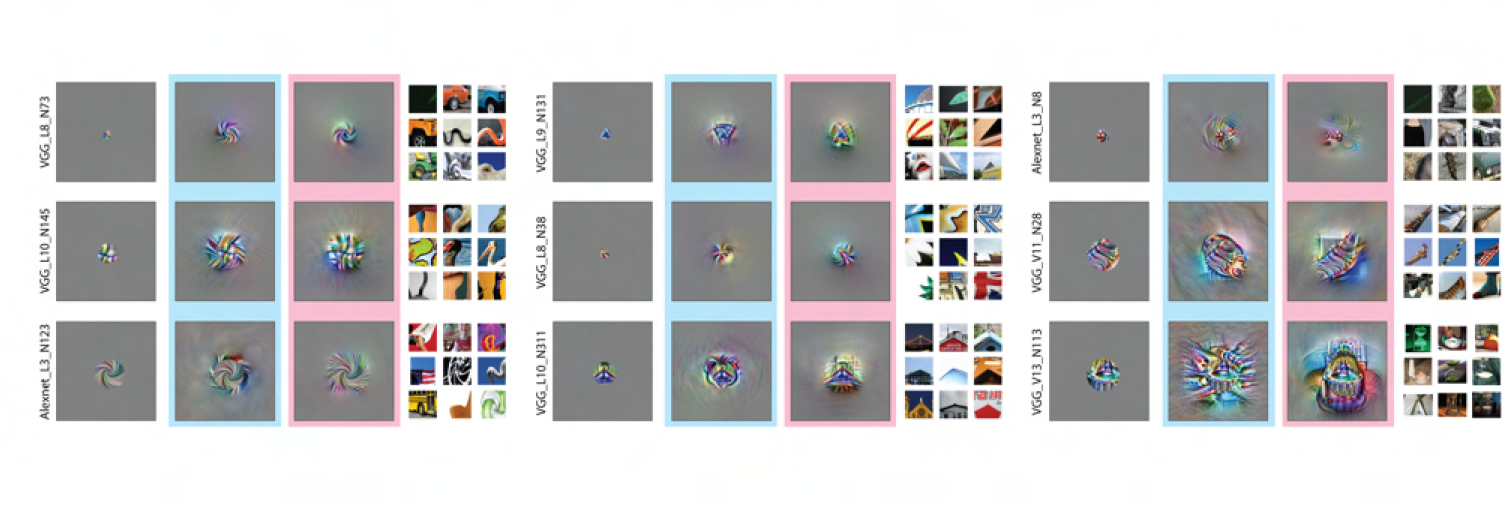
Examples of visualization of the most suppressive and facilitative surround for CNN neurons that do not show obvious homogeneity. The blue background denotes the most suppressive surround; the pink background denotes the most facilitative surround. The nine most excitatory natural image patches from the ImageNet validation set are shown on the right for each neuron. These neurons do not have clear center-surround contrastive features; they are likely to include surround features that are geometrically arranged rather than uniform across the surround or features that are arranged as object-like shapes.

**S9 Fig.**
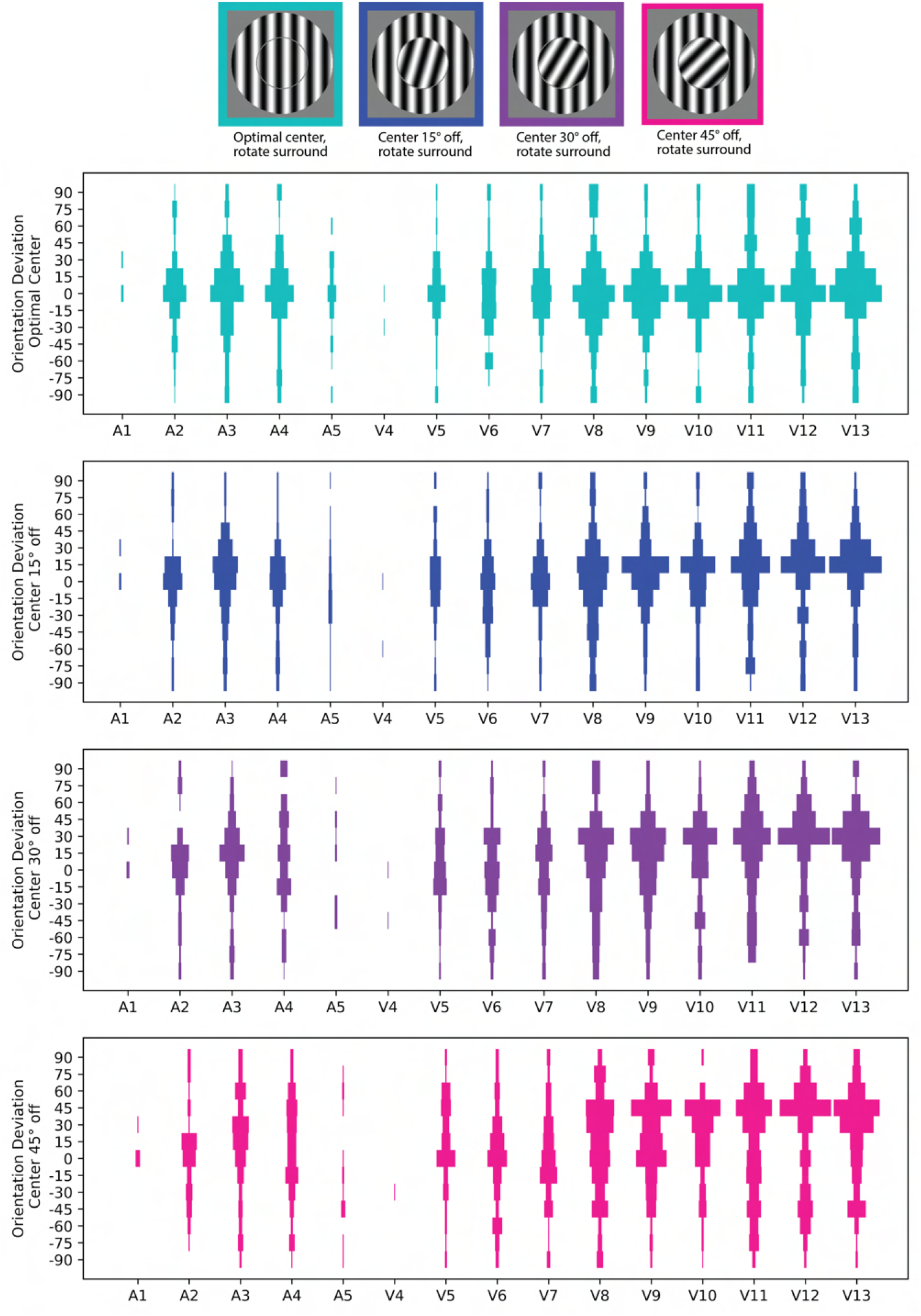
Histogram of surround suppression orientation deviation. Orientation deviation is defined as the difference between the most suppressive surround orientation and the optimal orientation. To plot this histogram, we included another selection criterion that the surround suppression tuning curve must have at least 0.001 variation, because if a surround suppression tuning curve is flat, there is not a meaningful orientation that has maximum suppression; therefore, it is not informative including in the histogram. In later layers (layer 8 and beyond) in VGG16, orientation deviations match the center orientation better on average.

**S10 Fig.**
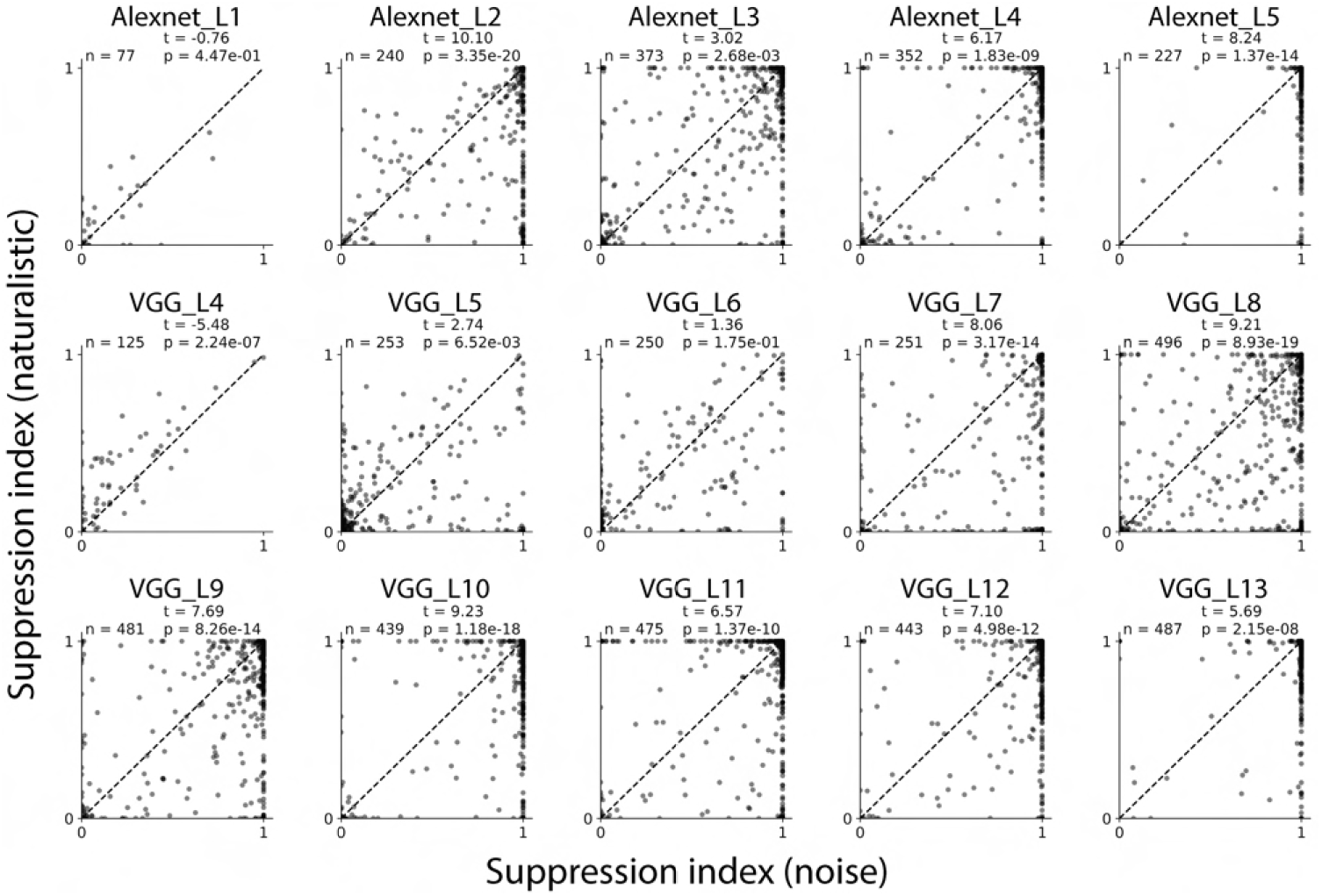
Naturalistic texture suppression index versus spectrally matched noise suppression index. Each dot represents a neuron. In most middle and later layers, neurons have higher suppression indexes with noise images than with naturalistic images, as indicated by the positive t-value and small p value. T and P values are from paired t-test.

**S11 Fig.**
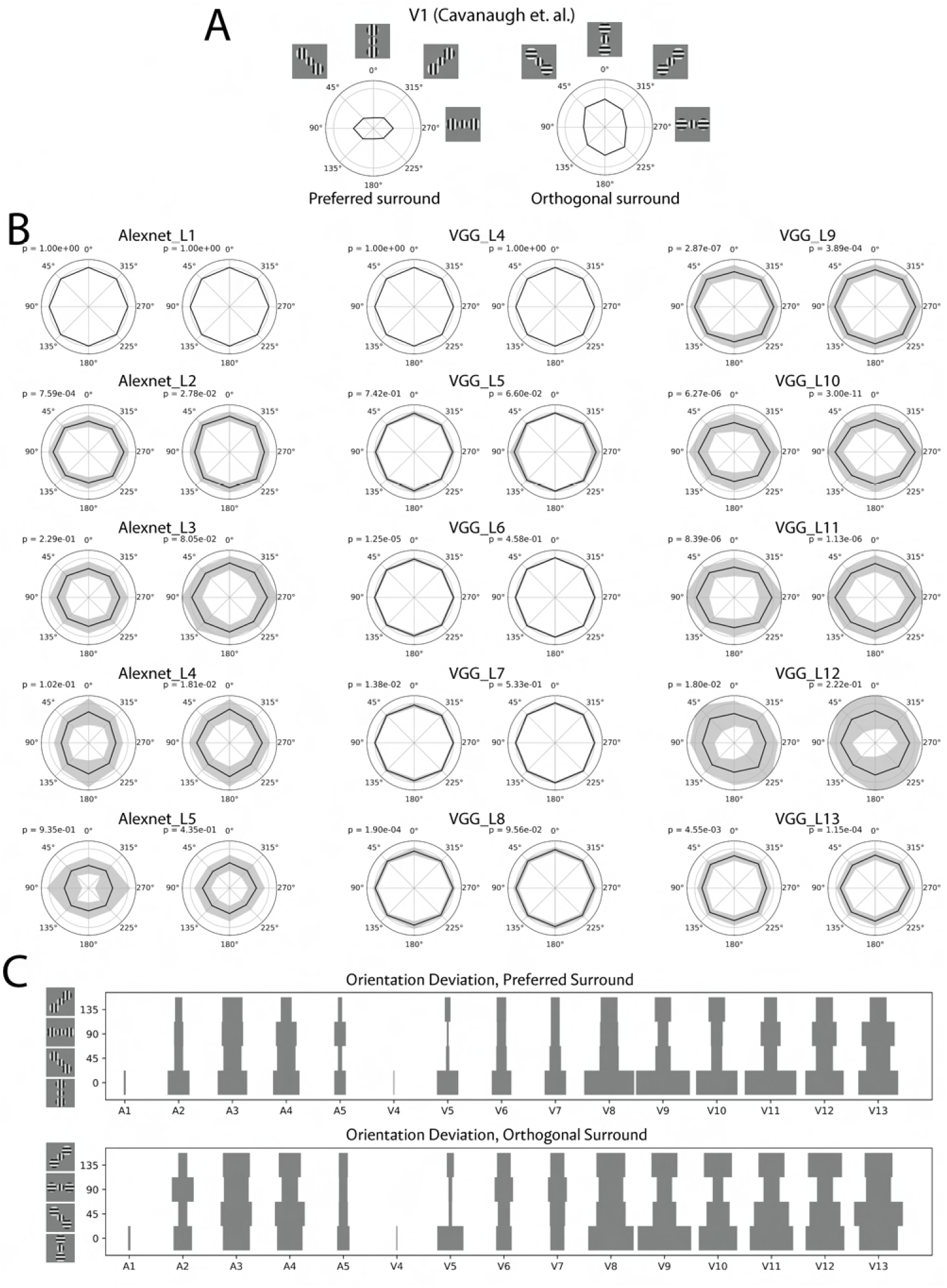
Geometry effects of the surround suppression. A. Strength of the surround suppression depends on the location of the surround stimuli. Embed images show the stimuli used in this experiment. The center was fixed at the optimal orientation. The surround had two patches at different locations relative to the center stimuli. The surround was either at optimal orientation or orthogonal orientation. Surround patches that align with the center stimuli induce the strongest suppression when it is at optimal orientation and the strongest facilitation when it is at orthogonal orientation. The polar radius represents the normalized response whereas the gray circle represents 1 (reproduced from [15]). B. Averaged plots of CNN layers. P values were calculated from one-way repeated measure ANOVA. Though some neurons and layers showed modulation effects of surround location, the effect size and shape of the plots did not match the cortical neurons shown in A. C. Histogram of the orientation deviation. Example stimuli of the four bins are shown on the left. When the surround is the preferred orientation, the most suppressive location is when they are collinear with grating orientation; when the surround is the orthogonal orientation, most layers still show the most suppressive when the surround location is collinear with the preferred orientation, which is inconsistent with electrophysiology data [15] where the most suppressive location is orthogonal to the optimal orientation.

**S12 Fig.**
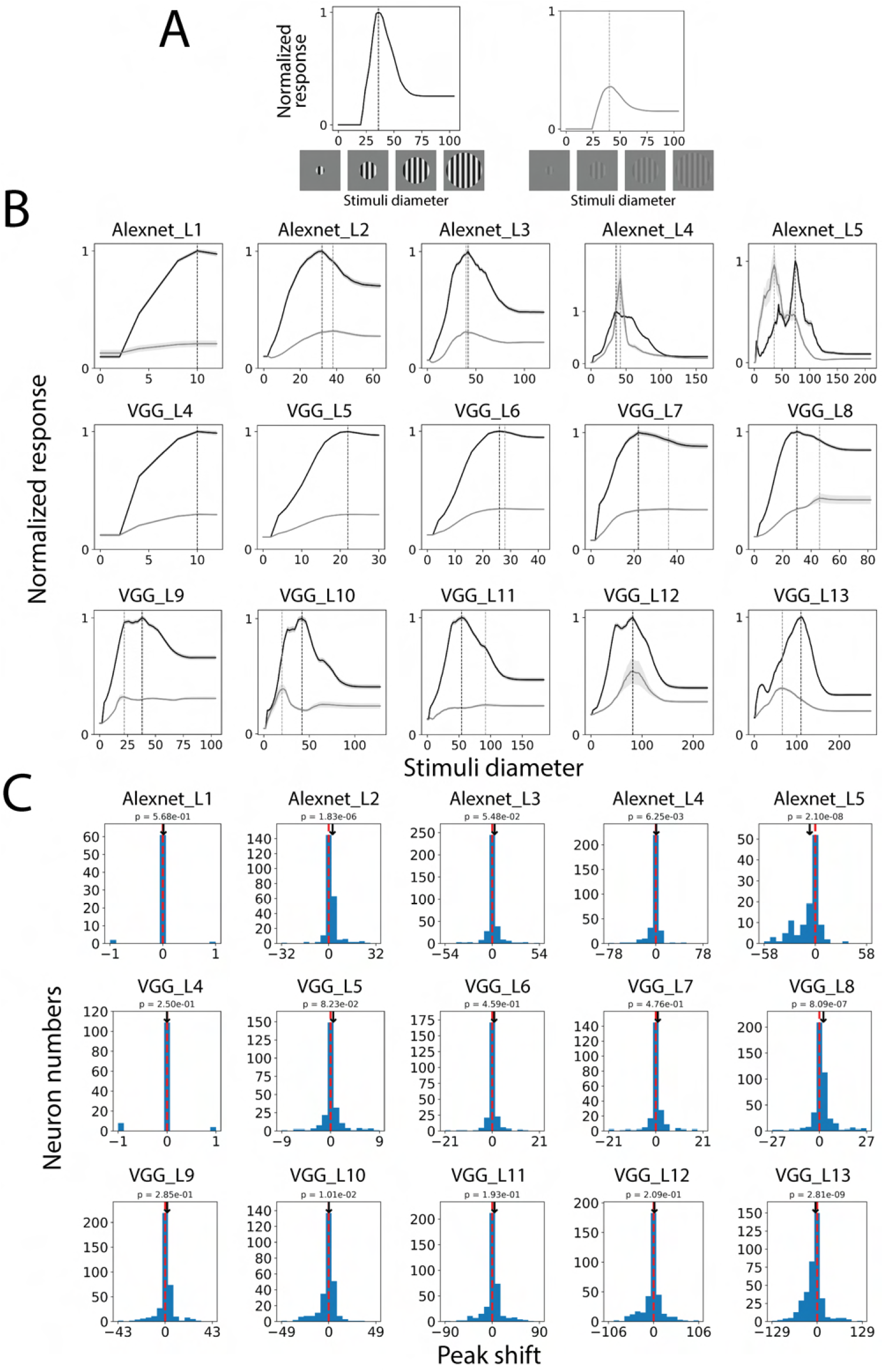
Low contrast does not consistently shift the peak of the diameter tuning curve as expected in neurophysiology. In neurophysiology studies, a low contrast grating causes the peak of the diameter tuning curves to shift to a larger size. We tested this effect in CNN neurons. A. Stimuli used in the simulation. Low contrast stimuli are at 17% of the regular pixel value range. B. Averaged diameter tuning curves with regular and low contrast stimuli. The black line denotes the regular contrast stimuli; the gray line denotes the low contrast stimuli. C. Histograms of the peak shift values. Positive values indicate low contrast stimuli shifted peak to a larger size, which is seen in cortical data. CNN neurons did not consistently show similar effects. In later layers, the low contrast peak even shifted to a smaller size significantly. Shaded area indicates s.e.m. P values are from paired t-test.

**S13 Fig.**
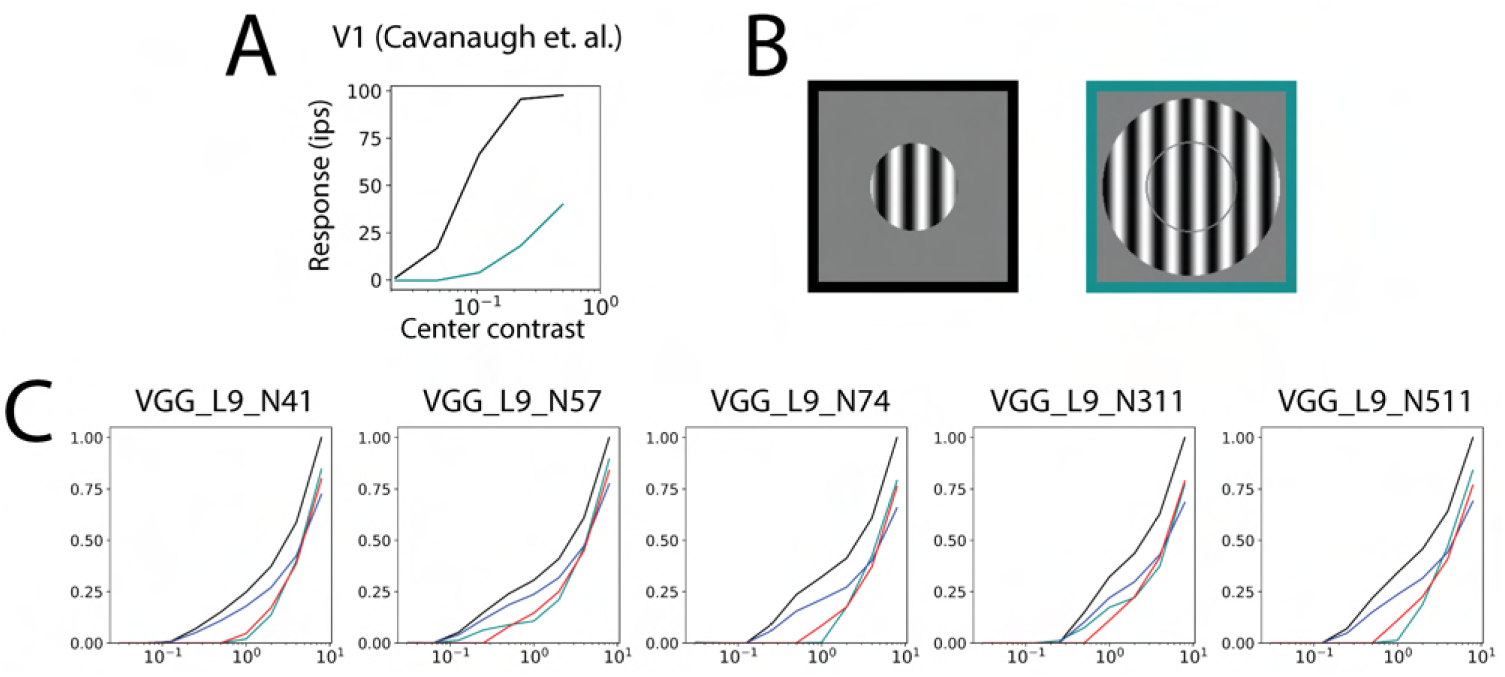
Center contrast responses for no surround and preferred surround. Contrast values were normalized to the regular pixel value range. We fixed the surround contrast at 1, changed the center contrast, and measured the contrast response function. We then fitted two curves with subtractive and divisive models. The subtractive model is described as Rs = max(0, Rc - a), where Rc is the responses of the center stimuli; Rc is the responses of the center stimuli with preferred surround; a is a subtractive parameter that is to be fitted. The divisive model is described as Rs = Rc/b, where b is a divisive parameter that is to be fitted. A. An example V1 neuron from a reference neurophysiology study (reproduced from [16]). Black line denotes no surround; cyan line denotes orthogonal surround. The contrast responses are shifted rightward and downward with surround suppression. B. Stimuli examples used in the experiments. C. Example CNN neurons with different behaviors. Blue line denotes a fitted surround suppression contrast curve with the divisive model; red line denotes a fitted surround suppression contrast curve with the subtractive model.

**S14 Fig.**
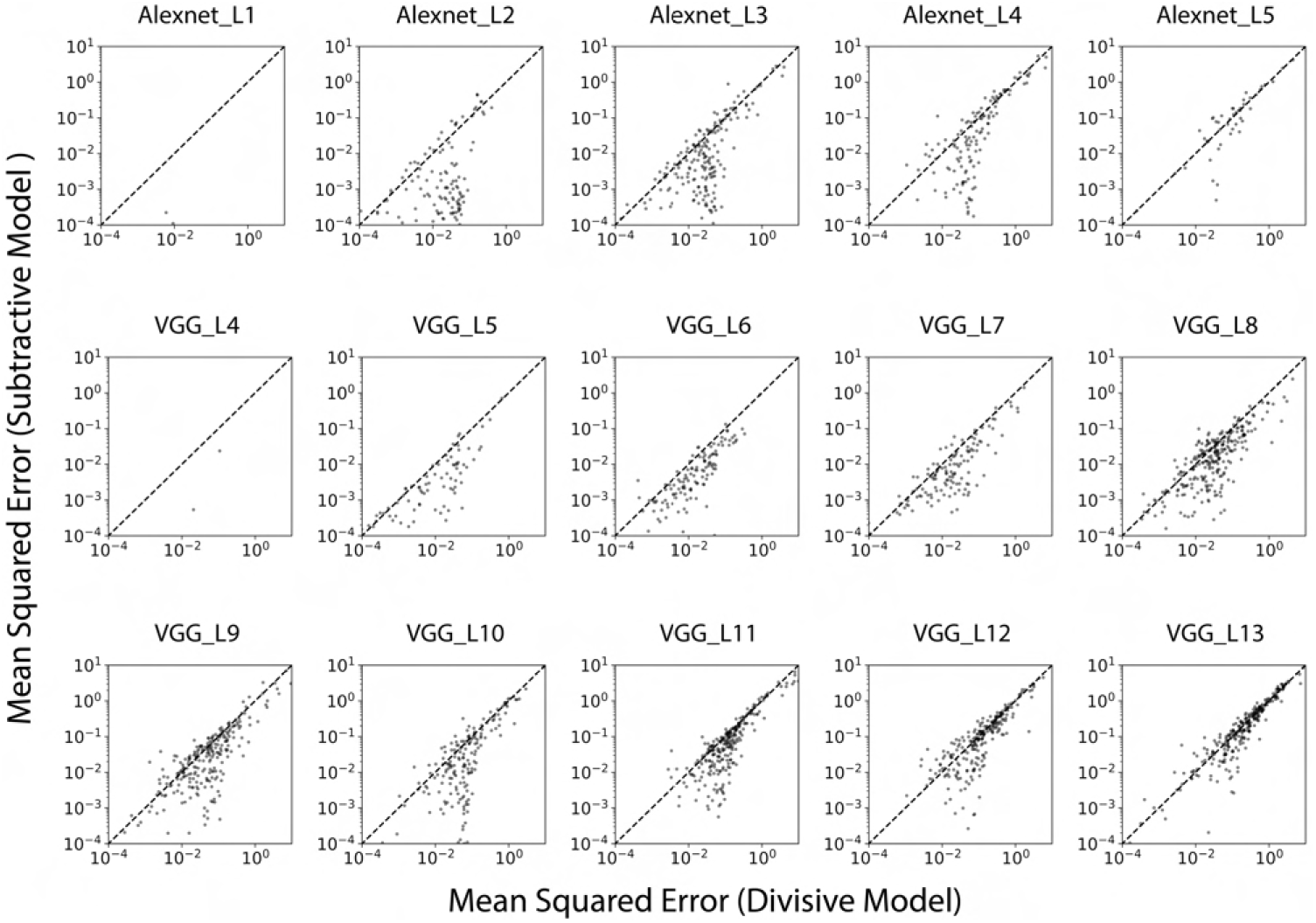
Explainability of subtractive and divisive models. To determine if the surround suppression effect is more likely to be in a subtractive form or divisive form, we fitted contrast curves in Supplementary Figure 9 with no surround and preferred surround by two models. The x-axis is the fitting error (mean squared error in log scale) of the divisive model; The y-axis is the fitting error of the subtractive model. Points below the diagonal line indicate neuron’s surround is more likely to be subtractive than divisive, which is commonly seen in most layers, especially early layers in both networks.

**S15 Fig.**
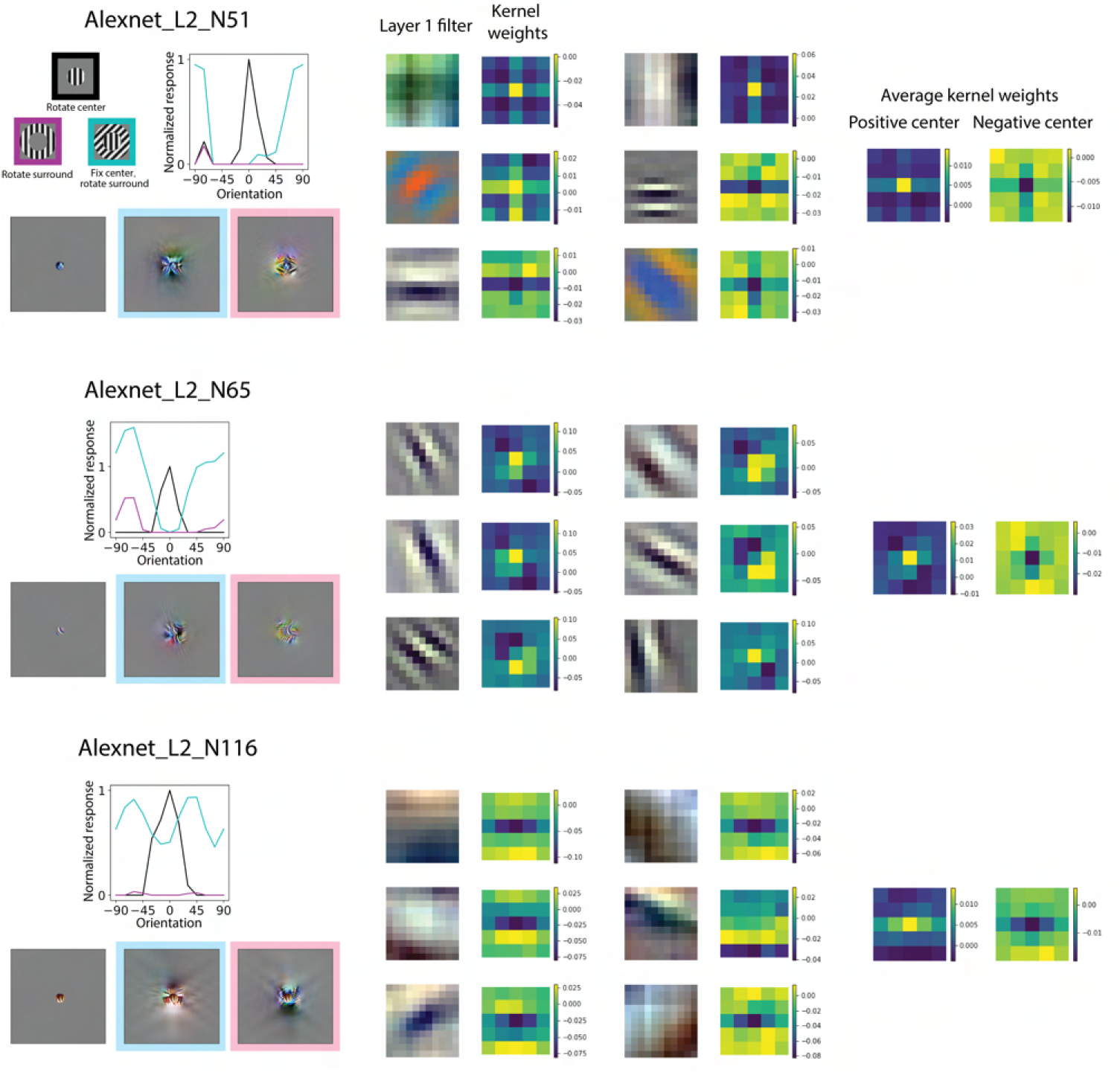
Surround suppression arises from the center-surround kernel weights. The homogeneous suppressive surround could be implemented by the center surround kernel weights structure. A subtractive form of surround suppression can be seen as the kernel weights having different signs in the center and surround. We show three example neurons in Alexnet layer 2. From left to right: tuning curves and visualization of the example neurons, six most contributing first-layer filters and corresponding kernel weights to the example neurons, and averaged kernel weights of kernels with positive or negative center weight. The most contributing first-layer filters are found by sorting the sum of absolute values of kernel weights. Most of the kernels have a center-surround structure. This indicates a subtractive form of surround suppression.

**S16 Fig.**
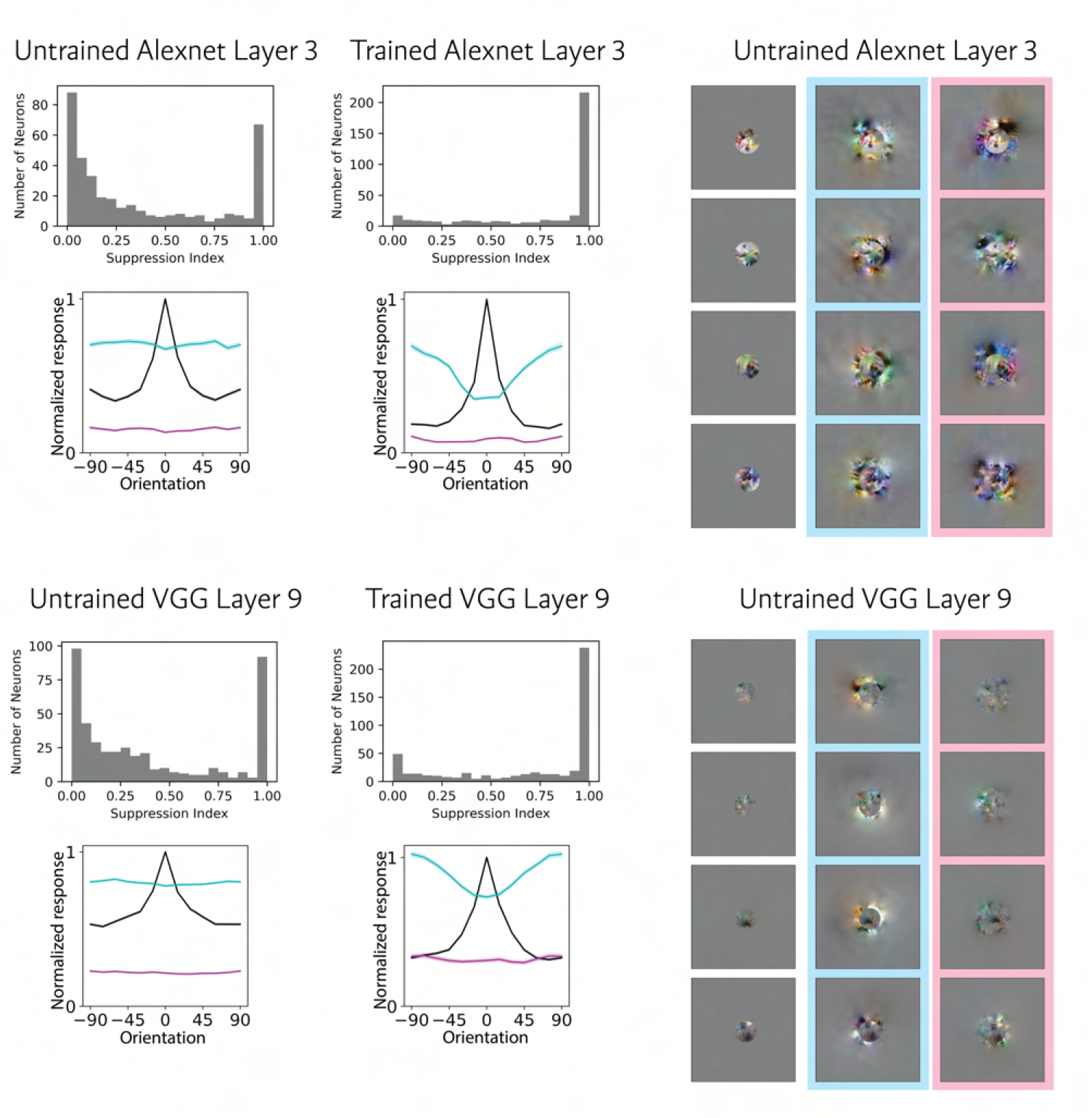
Untrained networks do not capture basic surround suppression effects. Suppression index, surround suppression curves and visualizations are shown for untrained Alexnet layer 3 and untrained VGG16 layer 9. While most neurons in trained networks showed close to 1 suppression index (strong surround suppression), neurons in untrained networks had bimodal distribution with the majority having close to 0 suppression index (no surround suppression). This was also reflected in the surround suppression curves (cyan), which in untrained networks were flat. On the right is the visualization of several example neurons in the untrained networks. There was no effects like most suppressive surround matches center. All the visualizations look like natural spectrum noise with no high-order textural appearance.

## Acknowledgments

We are grateful to Ruben Coen-Cagli for his advice and comments on the manuscript. This work was supported by a University of Miami Provost’s Research Award to O.S. A.D. was supported by the Research Experiences for Undergraduates (REU) Site *Scientific Computing for Structure in Big or Complex Datasets*, NSF grant CNS-1949972.

